# MicroRNA Tae-miR1130p targets wheat ferroportin1 (*TaFPN1*) in the absence of iron-responsive element/iron-regulatory protein1 module

**DOI:** 10.1101/2024.05.06.592689

**Authors:** Shivani Sharma, Riya Joon, Ajit Pal Singh, Deekshikha Tyagi, Gazaldeep Kaur, Ajay Kumar Pandey

**Author notes:** Centre for Genomic Regulation (CRG), The Barcelona Institute of Science and Technology, Dr. Aiguader 88, 08003 Barcelona, Spain.

## Abstract

Ferroportin (FPN) belongs to the Major Facilitator Superfamily of transporters and is a known iron (Fe) exporter in humans, with orthologues also present in plant species. Human FPN is subjected to multi-level regulation at transcriptional, post-transcriptional, and post-translational levels. How plant FPNs are regulated remains to be explored. In the current study, we have characterized wheat FPN1, a plasma-membrane localized protein, for its role in Fe homeostasis. A spatial-temporal expression analysis of wheat FPN1 suggested that its tissue-specific expression is differentially upregulated during Fe deficiency conditions. Unlike human FPN, plant FPN lacks the necessary sites for hepcidin binding, thereby emphasizing the need to explore the transcriptional/post-transcriptional mode of regulation. The lack of Iron Responsive Elements (IRE) in *TaFPN1* promoter suggests no direct regulation through the Iron Regulatory Protein (IRP) mechanism like in humans. Further, to explore the miRNA-mediated regulation, we identified Fe-regulated *tae-miR1130b-3p* capable of targeting *TaFPN1* under *in-vivo* conditions. Transcript expression of *tae-miR1130b-3p* negatively correlates with the *TaFPN1*. This alternative regulation pathway suggests a complex network of interactions governing the expression of genes involved in iron homeostasis, highlighting the intricacies of cellular regulatory mechanisms. Altogether, the work will unravel the cellular and physiological role of wheat FPN and contribute to a comprehensive understanding of plant iron homeostasis.

## Introduction

Iron (Fe) is an essential micronutrient with pivotal functions across all life forms. In plants, it is central to photosynthesis and chlorophyll biosynthesis, and a lack of it negatively impacts plant productivity and nutritional quality (Morrissey and Guerinot 2009; Connorton et al. 2017; Przybyla-Toscano et al. 2021). Being the fourth most prevalent metal in the earth’s crust, its solubility and availability vary depending on soil pH and Eh (redox potential) (Lindsay and Schwab 1982; Marschner 1995). Under alkaline and oxidative soil settings, Fe is rendered inadequate for uptake by plant roots due to the conversion of soluble Fe (II) into very insoluble Fe(III) oxyhydroxides. Thus, before plants can take up Fe for translocation into the root, these Fe(III) oxides must be solubilized. In contrast, Fe is readily available under acidic and submerged environments, resulting it to act as a potent toxin by causing oxidative damage through Fenton chemistry (Vose 1982; Hell and Stephan 2003). Therefore, a cell must constantly maintain iron in bound organic form because Fe^2+^ is prone to oxidation and precipitation inside the plant body and Fe^3+^ cannot be transported due to its low solubility (Olsen and Brown 1980; Lindsay 1991).

Land plants have evolved two main Fe acquisition strategies, referred to as Strategy I and Strategy II, to take sufficient Fe from the soil solution. All the dicots and non-graminaceous monocots utilize Strategy I, whereas Strategy II is employed by the Gramineae (Römheld and Schaaf 2004; Conte and Walker 2011; Kobayashi and Nishizawa 2012). Fe acquisition in Strategy I plants is mediated by a tripartite protein complex that causes apoplast acidification, reductive splitting of Fe^3+-^chelates, and uptake of the liberated Fe^2+^ ion. This complex comprises the H^+^ -ATPase AHA2, the oxidoreductase FRO2, and the Fe^2+^ transporter IRT1. Grasses (such as wheat, rice or barley) follow strategy II to mobilize and absorb Fe using phytosiderophores; high-affinity metal chelators of the mugineic acid family. TOM1 secretes phytosiderophores, followed by the transport of the loaded Fe(III)-PS complexes by high-affinity YS1 and YSL transporters (Curie et al. 2001; Inoue et al. 2009; Nozoye et al. 2011). Rice and potentially other grasses, like the Strategy I plant, have IRT1 homologs that aid in the uptake of (ferrous) Fe when Fe^2+^ is high, such as under waterlogged conditions (Morrissey and Guerinot 2009; Ishimaru et al. 2010; Zanin et al. 2017; Kaur et al. 2019)

Fe homeostasis in plants revolves around its efficient uptake, transport, and storage, with all these processes tightly regulated (Walker and Connolly 2008; Hindt and Guerinot 2012). In addition to iron acquisition and compartmentalization, identifying the transporters that govern Fe transit from the root to the shoot is crucial for properly understanding iron homeostasis. Genetic dissection of Fe transporters and regulatory checkpoints have been addressed in dicot and monocot plants such as *Arabidopsis thaliana, Zea mays, Hordeum vulgare and Brachypodium distachyon* (Murata et al. 2006; Bashir et al. 2010; Yordem et al. 2011; Gómez-Galera et al. 2012; Brumbarova et al. 2015; Zhang et al. 2022). But in case of hexaploid wheat, gene mining has been difficult until recently owing to its complicated genomic architecture. Nonetheless, high throughput omics approaches have contributed to identifying Fe deficiency responsive novel Strategy II players required for Fe loading from the root to the shoot (Wang et al. 2019; Kaur et al. 2019).

After the uptake of Fe through IRT1 by root epidermal cells, it travels symplastically through the root cortex, the endodermis - including the Casparian strip, and the pericycle where it is released in the vasculature through the Ferroportin 1 (FPN1). Ferroportin (FPN), also known as iron-regulated transporter 1 (IREG1) belongs to the solute carrier family 40 (SLC40A1) of the Major Facilitator Superfamily (MFS), is the sole known iron exporter in vertebrates localised to the membranes of enterocytes, macrophages hepatocytes and adipocytes (McKie et al. 2000; Donovan et al. 2005; Drakesmith et al. 2015). FPN regulates iron release into the plasma and is also a door to iron recycling inside the macrophages (Abboud and Haile 2000; Hentze et al. 2004). Lately, the vertebrate *FPN* homologs have also been addressed in plant systems like *Arabidopsis thaliana*, *Oryza sativa*, *Fagopyrum esculentum*, *Psychotria gabriellae* and *Medicago truncatula* (Morrissey et al. 2009; Merlot et al. 2014; Yokosho et al. 2016; Escudero et al. 2020; Kim et al. 2021; Kan et al. 2022). Plasma membrane-localized exporter known as Ferroportin 1 (FPN1) was identified in *Arabidopsis thaliana* (Morrissey et al. 2009). AtFPN1 was found to be expressed in the root and shoot vasculature and performed the loading of Fe and cobalt (Co) from the pericycle to the xylem. Besides *AtFPN1*, two other *FPN/IREG* family members i.e., *AtFPN2, and AtFPN3* were also discovered in *Arabidopsis.* AtFPN2 localized on the tonoplast and expressed in the root epidermis and cortex, where it performed sequestration of Co and nickel (Ni) into vacuoles for detoxification (Schaaf et al. 2006; Morrissey et al. 2009). AtFPN3 was dual targeted to the chloroplasts and mitochondria. (Conte et al. 2009; Kim et al. 2021). Recently, multiple *FPNs* were identified in *Medicago truncatula* referred to as *MtFPN1, MtFPN2, and MtFPN3*.

Functionally characterized *MtFPN2* expressed in the nodule vasculature and performed iron transport into the nodules, implying a function in symbiotic nitrogen fixation (Escudero et al. 2020). *FPN1* of *Psychotria gabriellae* (*PgFPN1*) and buckwheat/*Fagopyrum esculentum Moench* (*FeFPN1*) played specifically crucial roles in detoxification of Ni and Al^3+^, confirmed by their overexpression studies in *A. thaliana* (Merlot et al. 2014; Yokosho et al. 2016). In rice, *OsFPN1* was localized on the golgi apparatus and plasma membrane and performed Co and Ni detoxification in roots.(Kan et al. 2022).

Regulation of FPN is quite unique and most of the knowledge is derived from its detailed characterization in mammalian system. Human FPN is regulated at multiple levels, including transcription, post-transcriptional, and post-translational. The *FPN* promoter contains HIF-responsive elements (HRE) as well as metal response elements for zinc, manganese, and copper, implying that it is regulated by hypoxia and transition metals (Troadec et al. 2010; Taylor et al. 2011; Mitchell et al. 2014). Stem-loop iron-responsive element (IRE) motifs are also present in the *FPN* promoter. When iron levels are low, iron regulatory proteins (IRPs) bind to IREs in the 5’ untranslated region (UTR) of transcripts, where they impede translation, or in the 3’ UTR of transcripts, where they hinder degradation when Fe is replete (Zhang et al. 2009; Zhou and Tan 2017). In another mode of FPN regulation, microRNA-mediated post-transcriptional regulation of FPN has been demonstrated by targeting its UTR region. Human miR-485-3p has been shown to bind to the 3’ UTR of FPN and suppress its translation (Sangokoya et al. 2013). This miRNA was found to be downregulated during high Fe levels were high, thus resulting in enhanced FPN expression. FPN is also regulated post-translationally by hepcidin, an amphipathic peptide of 25 amino acids that facilitates FPN internalization and destruction in lysosomes (Nemeth et al. 2004; Ross et al. 2012). Despite progress in understanding Fe transport by FPN and its regulation in humans,there is a large gap in the knowledge of the regulation of plant *FPN1* via the above three conserved mechanisms.

Hexaploid wheat (*Triticum aestivum* L.) is the most widely cultivated crop globally and is responsible for fulfilling the nutritional needs of one-fifth of the population of the world. The functional role of FPN and its mode of regulation still need to be explored in plants. Therefore, it is imperative to investigate the roles of Fe transporters primarily involved in loading iron from root to shoot to yield an in-depth understanding of micronutrient homeostasis. In the current work, we provide functional evidence that shows complementation of the wheat FPN1 in *atfpn1* mutant plants and regulation of *TaFPN1* transcript primarily mediated by miRNA.

## Materials and Methods

### Plant Material and growth conditions

Hexaploid bread wheat (*Triticum aestivum*) cv. “C306” was used for expression analysis. To analyze gene expression in growing seeds, the main individual spikes of three biological replicates were tagged the next day after anthesis (DAA) and harvested at four developmental stages, i.e., 7, 14, 21, 28 DAA. Tissue specific expression analysis was conducted in root, stem, leaf, flag-leaf, and seed samples collected at 14DAA. To study the response of *TaFPN1* under different metal stress regimes, its temporal expression was studied in roots and shoots of C306 wheat plants grown in hydroponic setup. To give Fe treatments, seeds of *Triticum aestivum* cv. C-306 were washed with water and sterilized with 1.5% sodium hypochlorite.

They were allowed to germinate at room temperature and then healthy seedlings were grown in Hoagland’s nutrient solution in the growth chamber for 5 days. For Fe deficiency (–Fe), 2 μM Fe (III) EDTA and for Fe surplus (++Fe) conditions 200μM Fe (III) EDTA was used as the Fe source. For control plants (+Fe), concentrations of nutrients were unchanged and kept at 80 μM Fe (III) EDTA. Plants were grown at 21±1 °C, 50–65 % relative humidity, and a photon rate of 300μmol quanta m^−2^ s^−1^ with a 16 h day/8 h night cycle. For sampling, roots and shoots were collected after 3, 6, 9, 12 and 15 days of Fe stress treatment from three biological replicates. Furthermore to understand the responsiveness and specificity of *TaFPN1* towards heavy metals, its expression profiling was performed in the root and shoot tissues of wheat plants subjected to Cadmium stress (+Cd; 50 µM), Cobalt stress (+Co; 50 µM) and Nickel stress (+Ni; 50 µM) in Hoagland media. For functional characterization of wheat iron transporter (*TaFPN1*) in-planta, the Col-0 line *fpn1-2* (SALK_145458) was employed.

### In-silico identification of wheat FPN genes and Phylogenetic analysis

To obtain wheat ferroportin genes (*TaFPNs*) from the whole genome of wheat, BLAST program (https://plants.ensembl.org/Triticum_aestivum/Tools/Blast) was employed for identification of *TaFPN* homologs and two online websites, Pfam v35.0 (Protein Family database) (http://Pfam.xfam.org) and NCBI-CDD (National Center for Biotechnology Information-Conserved domain database), were used for confirmation. The FPN protein sequences of *Arabidopsis thaliana* (AtFPN1, AtFPN2 and AtFPN3) and *Oryza sativa* (OsFPN1, OsFPN2 and OsFPN3) were obtained from relevant literature reports (Morrissey et al. 2009; Kim et al. 2021; Kan et al. 2022) and species specific databases; The Arabidopsis Information Resource (TAIR) (https://www.arabidopsis.org/) and The Rice Annotation Project Database (RAP-DB) (https://rapdb.dna.affrc.go.jp/) (Rhee et al. 2003; Sakai et al. 2013). The obtained protein sequences were employed as reference sequences to identify potential *TaFPN* genes by performing BLAST P searches with an e-value <1e^−10^ against the wheat genome assembly (IWGSC RefSeq v.1.1) of Ensembl Plants Database (https://plants.ensembl.org/index.html).

To classify wheat FPNs a phylogenetic tree was constructed with wheat FPN protein sequences along with FPN protein sequences from *Arabidopsis thaliana* (AtFPN1), AT5G03570 (AtFPN2), AT5G26820 (AtFPN3), *Oryza sativa*; Os06t0560000 (OsFPN1), Os12t0562100 (OsFPN2), Os05g0131500 (OsFPN3), *Medicago truncatula*; Medtr1g084140 (MtFPN1), Medtr4g013240 (MtFPN2), Medtr4g013245 (MtFPN3), *Homo sapiens* (NP_055400.1), *Mus musculus* (NP_058613.2) and *Bdellovibrio bacteriovorus* (WP_011164469.1). FPNs from all species were aligned through MUSCLE algorithm and then subjected to construct an unrooted tree using neighbour-joining method with 1000 bootstrap value using MEGA 11 software.

### RNA extraction and gene expression analyses

Total RNA extractions were performed using Trizol Reagent, as described earlier (Rio et al. 2010). The quantity and purity of RNA was examined using NanoDrop™ Lite Spectrophotometer (Thermofisher Scientific). The contamination of genomic DNA was removed by subjecting the RNA samples to DNase treatment using TURBO™ DNase (Ambion, Life Technologies). For qRT-PCR, cDNA was synthesized from DNase-free total RNA using Invitrogen SuperScript III First-Strand Synthesis System SuperMix (ThermoFisher Scientific). Gene expression was quantified using the QuantiTect SYBR Green RT-PCR Kit (Qiagen, Germany) in Bio-Rad CFX96 Real-time PCR detection system. The gene-specific primers capable of amplifying 150-250 bp region from all the three homoeologs of *TaFPN1* gene were carefully designed using Oligocalc software. Primers used in qRT-PCR are illustrated in (Table S1).The relative transcript accumulation was calculated following the 2^−ΔΔCT^ method (Livak and Schmittgen 2001), using ADP-ribosylation factor gene (*TaARF*) as an internal control.

### Quantification of miRNAs by Stem-loop qRT-PCR

To get an insight into the spatio-temporal expression patterns of differentially expressed miRNAs targeting *TaFPN1*, stem-loop qRT-PCR analysis was conducted. Primers were designed as described in the literature reports (Chen et al. 2005). Briefly, 1μg of DNase-treated total RNA was reverse-transcribed using TaqMan microRNA reverse transcription kit (Applied Biosystems^TM^) according to the manufacturer’s instructions. The real-time PCR program was set as follows: 95°C for 3 min, 45 cycles (95°C for 10 sec, 55°C for 20 sec,72°C for 20 sec). All reactions were performed in triplicate for each time point. The relative expression levels of the miRNAs were calculated by the 2^-ΔΔCT^ method (Livak and Schmittgen 2001). For normalization, the wheat *U6* snRNA gene (GenBank: X63066.1) was used as an internal control (Gasparis et al. 2017). All qRT-PCRs was performed using SYBR Green I (Takara SYBR Premix Ex Taq) on the Bio-Rad CFX96 Real-time PCR detection system.

### Vector construction and plant transformation

A full-length sequence of *TaFPN1* was amplified using pooled cDNA prepared from 1-2 μg of total RNA, extracted from the roots of 9-15 days old C306 wheat cultivar subjected to Fe deficiency (2 μM Fe-III EDTA). Primers were designed from the conserved region of *TaFPN1* homoeologues (Table S1). Phusion High-Fidelity DNA Polymerase (ThermoFisher Scientific) was used for PCR amplification and the amplified gene was cloned in pJET1.2 vector (ThermoFisher Scientific). The sequence was confirmed using 3730xl DNA Analyzer (Applied Biosystems).

To prepare the overexpression construct for the transformation of Arabidopsis *fpn1-2* (SALK_145458) mutants, *TaFPN1* was cloned in Gateway-compatible plant expression vector - pGWB520. Firstly, an entry clone of *TaFPN1* was generated by sub-cloning the gene in the pENTR™/D-TOPO™ vector followed by recombination of the *TaFPN1* gene fragment in the destination vector;pGWB520 using LR clonase II enzyme mix (Thermofisher Scientific). The resulting construct; *TaFPN1*-pGWB520 was then used to transform *Agrobacterium* GV3101 strain using the floral dip method (Clough and Bent, 1998).

Transformants were isolated by plating the seeds on half MS media with hygromycin (25 mg/mL). Homozygous T3 lines showing a clear 100% resistance to hygromycin B were used for further quantitative analysis.

### Sub-cellular localization of TaFPN1

*TaFPN1* amplification was performed from the sequence confirmed positive *TaFPN1-* pJET1.2 clone. Primers were designed using the Neb Hi-fi builder assembly tool (https://nebuilder.neb.com) by adding *Nco*I restriction enzyme site in the forward primer and *SpeI* in the reverse primer respectively. The *TaFPN1* fragment was ligated in the destination vector pCAMBIA1302 using Gibson assembly master mix (Replica express kit, Genes2Me). Consequently, the *TaFPN1-GFP* construct, pCAMBIA1302 empty vector, and plasma membrane marker clone pMRK-RFP were mobilized into Agrobacterium strain GV3101. This was followed by resuspending the transformed GV3101 cells in agroinfiltration medium [10 mM MES (pH 5.6), 10 mM MgCl_2_, and 200µM acetosyringone] at an OD_600_ of 0.6-0.8 and inoculating the abaxial surfaces of six-eight week old *Nicotiana benthamiana* leaves using syringe infiltration. The leaf tissues were visualized for fluorescent signal 48h post infiltration using LEICA DM600CS confocal microscope. GFP fluorescence was detected at 500–530 nm, whereas the red fluorescence was detected at 580–630 nm.

### Chlorophyll estimation and measurement of Fe content

For the assessment of shoot chlorophyll concentrations, plants were grown on B5 medium for 5 days at 22 ± 1L, 16 h light/8 h dark cycle and a photon rate of 100µmol photons m^-2^ s^-1^ and then transferred to minimal media that was either iron sufficient (Fe (III) EDTA, 50µM), deficient (–Fe media supplemented with 300µM of Ferrozine) or surplus (Fe (III) EDTA, 200µM). The shoots were harvested and subjected to chlorophyll extraction in 80% acetone containing 20% (v/v) 0.2M Tris–HCl, pH 8. The leaves were incubated at room temperature in a 1.5-mL tube with 1 mL 80% acetone solution for at least 24 h and was clarified by centrifugation for 5 min at 15,000g. The absorbance of the supernatant was measured at wavelengths 646 and 663 nm (A646 and A663) by spectrophotometry. Chlorophyll concentration was estimated using the Lichtenthaler’s equation (Lichtenthaler 1987) :

Total Chlorophyll (µg/mL) = 6.43A_663_ + 18.43A_646_ Fe estimation was done as described earlier (Meena et al., 2024). For total Fe analysis, *TaFPN1* overexpressing *Arabidopsis* lines, *fpn1-2* mutant and wild type Col-0 plants were grown in B5 media for five days and then transferred to minimal B5 media supplemented with +Fe (50µM) for two weeks and analysed for Fe distribution by ICP-MS. The tissue was finely ground and dried at 60 °C to yield 10 mg of tissue which was then digested using 2.5mL concentrated HNO_3_ in a microwave reaction system (Mars 6, CEM Corporation, USA) and diluted to 10 mL with 5% HNO3. The digested samples were subjected to total Fe analysis using inductively coupled plasma mass spectrometry (ICP-MS; 77006 AgilentTechnologies, Santa Clara,CA). To ensure the accuracy of the analysis, two technical replicates were performed for each of the three biological replicates. The metal content was measured by normalizing the sample metal concentration (ppb) with the dry mass of respective plant sample. To determine the Fe distribution pattern, the *Arabidopsis* seedlings were germinated on Gamborg’s B5 agar media for 5 days and then shifted to B5 minimal media amended with +Fe (control), −Fe (Fe-deficient) and ++Fe (Fe surplus) for 10 days.

The plant samples were washed with distilled water. Subsequently, they were vacuum infiltrated in a solution of 4% HCl and 4% K-ferrocyanide for 30 minutes. Following this, a thorough washing with distilled water was done to remove any excess staining solution.

Furthermore, the plants were kept in a freshly prepared solution composed of 0.1% DAB (3,3’-Diaminobenzidine) in 0.1 M Sodium citrate buffer, pH 3.8 under dark conditions. A final rinse with distilled water was done to remove any residual dye. Finally, different stained plant regions were examined under a stereo microscope (Leica Microsystems, Germany).

### Screening of IRE elements in wheat genome and interaction with IRP

The predicted wheat genes containing the iron responsive elements (IRE) in either 5’ or 3’ UTR. CxxxxxCAGUGNyyyyy IRE sequence was looked for in all UTRs (x and y complementary as AU/UA/GC/CG/UG/GU using an in-house generated Perl script. Further, we used SIREs (Searching for Iron-Responsive Elements) (http://www.sires-webserver.eu/), the bioinformatic program for the prediction of IREs. All motifs were checked for the characteristic C-bulge (C8) present in the stem motif and a 6-nucleotide – CAGAGU/C- apical loop, both circled in Red. The standard IRE hairpin-loop structure consists of a six-nucleotide loop with specific sequences (5’-CAGWGH-3’, where W represents A or U and H represents A, C, or U) connected to a stem comprising five paired nucleotides. Additionally, a slight asymmetrical bulge contains an unpaired cytosine on the 5’ side of the stem and a variable-length lower stem.

Human IRP and rice aconitase (*OsACO*) were cloned in the pACT2-AD vector, and wheat IREs and human IRE were cloned in the pIIIA/MS2-2 vector (Stumpf et al. 2008) (Gifted by Prof. Marvin P. Wickens). After being co-transformed in the YBZ-1 yeast strain, the produced constructs were plated on SD/-Leu/-Ura. For interaction experiments, the co-transformed cells were also plated on SD/-Leu/-Ura/-His plates. Pictures were taken after 4 days of growth.

### Construct designing for tae-miR1130b-3p and TaFPN1

The *TaFPN1* sequence region encompassing the 21 bp target site for the *tae-miR1130b- 3p* was used to create forward and reverse primers. These target site primers were then subjected to form a duplex utilizing PCR with a gradient reduction of 0.1°C/cycle. The duplex so formed was ligated and fused to *GUS* gene in pCAMBIA1301 vector. For cloning of miRNA, the pre-miRNA sequence of *tae-miR1130b-3p* was employed to design forward and reverse primers consisting of 18-20 nucleotide stretch of overlapping sequence.

Subsequently, overlap extension PCR was employed to generate *tae-miR1130b-3p* duplex followed by cloning in pENTR/D-TOPO entry vector and subsequently in the destination vector - pMDC32. The constructs were then mobilised into *Agrobacterium* strain GV3101 and six-eight week-old *Nicotiana benthamiana* plants were agro-infiltrated at an OD of 0.6-0.8. Half of the abaxial surface from the mid-rib was infiltrated with target site construct and half with the target site + miRNA construct.

### Quantification of GUS enzyme activity

Fluorometric quantification of GUS enzyme activity was performed as described previously (Blázquez 2007) with minor modifications. Leaf samples were crushed in LN_2_ followed by the addition of Sodium Phosphate buffer (pH 7.0) and total protein was extracted. Total protein concentration was determined with a Bradford Protein Assay Kit (Takara Bio Inc.) according to the microtiter plate protocol recommended by the manufacturer. For MUG (4- methylumbelliferyl- -D-glucoside) assay reaction, 5µg of protein was mixed with 130µl of MUG assay buffer and incubated at 37°C for 30 min. The reaction was terminated by adding sodium carbonate (stop buffer). Quantification of 4-MU was performed at excitation at 355 nm and emission at 465 nm. GUS activities were expressed as fluorescence intensity per µg of total protein.

## Results

### Wheat genome encodes multiple Ferroportins (FPN1, FPN2 and FPN3)

In an attempt to identify the wheat *FPNs*, multiple *Arabidopsis* and *Oryza sativa* FPNs protein sequences were used to perform the homology searches in the Wheat Ensembl database. This resulted in identifying 9 wheat *FPN* homoeologues belonging to the Solute carrier member family 40. Analysis of the TaFPN protein sequences in the Pfam database revealed that they all possessed a functional domain which is the characteristic of the Major Facilitator Superfamily (MFS) of transporters (pfam; PF06963). Using phylogeny analysis, they were categorized as *TaFPN1*, *TaFPN2* and *TaFPN3* on the basis of the degree of shared identity with other plant *FPN* orthologues (Figure 1A & Figure S1). TaFPN1 & TaFPN2 homoeologues shared >80% sequence identity with each other at the protein level, >74% sequence identity with OsFPN1 and >54% identity with AtFPN1 and AtFPN2. In comparison, TaFPN3 shared only 23-25% identity with TaFPN1 and TaFPN2, but >80% identity with OsFPN2 and >52% identity with AtFPN3. Interestingly, all the wheat FPN homoeologs were present on the A, B, and D sub-genomes, suggesting all the wheat progenitors contained *FPN* encoding sequences. Earlier research has demonstrated the crucial roles of plant FPNs during Fe homeostasis (Morrissey et al. 2009; Kim et al. 2021; Kan et al. 2022) . Therefore, we checked the expression of the wheat *FPNs* in the previously generated transcriptomic datasets derived from wheat C306 roots subjected to Fe deficiency condition (Kaur et al. 2019; Kaur et al. 2023) . To check which wheat *FPN* responds to Fe deficiency, a heat map was generated for all the identified wheat *FPNs*. Interestingly, our analysis revealed significant induction of all the homoeologs of the *TaFPN1* gene at multiple time points of Fe deficiency (Figure 1B). None of the *TaFPN2/TaFPN3* homologs showed any expression response in the dataset. Subsequently, we validated the temporal expression response of *TaFPN1* under Fe deficiency and surplus conditions in root and shoot tissues. qRT-PCR expression analysis of *TaFPN1* revealed its significant upregulation in response to Fe deficiency in the roots and shoots at both early and late time points. Under Fe surplus conditions, *TaFPN1* showed downregulation in the roots and shoots, specifically during late time points (Figure 1C & 1D). Expression of the *TaFPN1* was induced at all the time points studied, suggesting its response to Fe deficiency conditions. Specifically, the highest expression was observed at 15 days of Fe deficiency in roots. (Figure 1C). Subsequently, *TaFPN1* was assessed for localization in the leaf epidermal cells of *N. benthamiana* by transiently co-expressing TaFPN1-GFP with the plasma membrane marker pMRK-RFP. As a result, the TaFPN1-GFP fusion protein was observed at the plasma membrane. This agreed well with the FPN1 localization of *Arabidopsis thaliana* and *Oryza sativa,* which were also found to localize on the plasma membrane (Morrissey et al. 2009; Kan et al. 2022). (Figure 1D). Overall, our data indicated that the *TaFPN1* might function as a potential plasma membrane-localized Fe efflux during Fe homeostasis.

**Figure 1:**
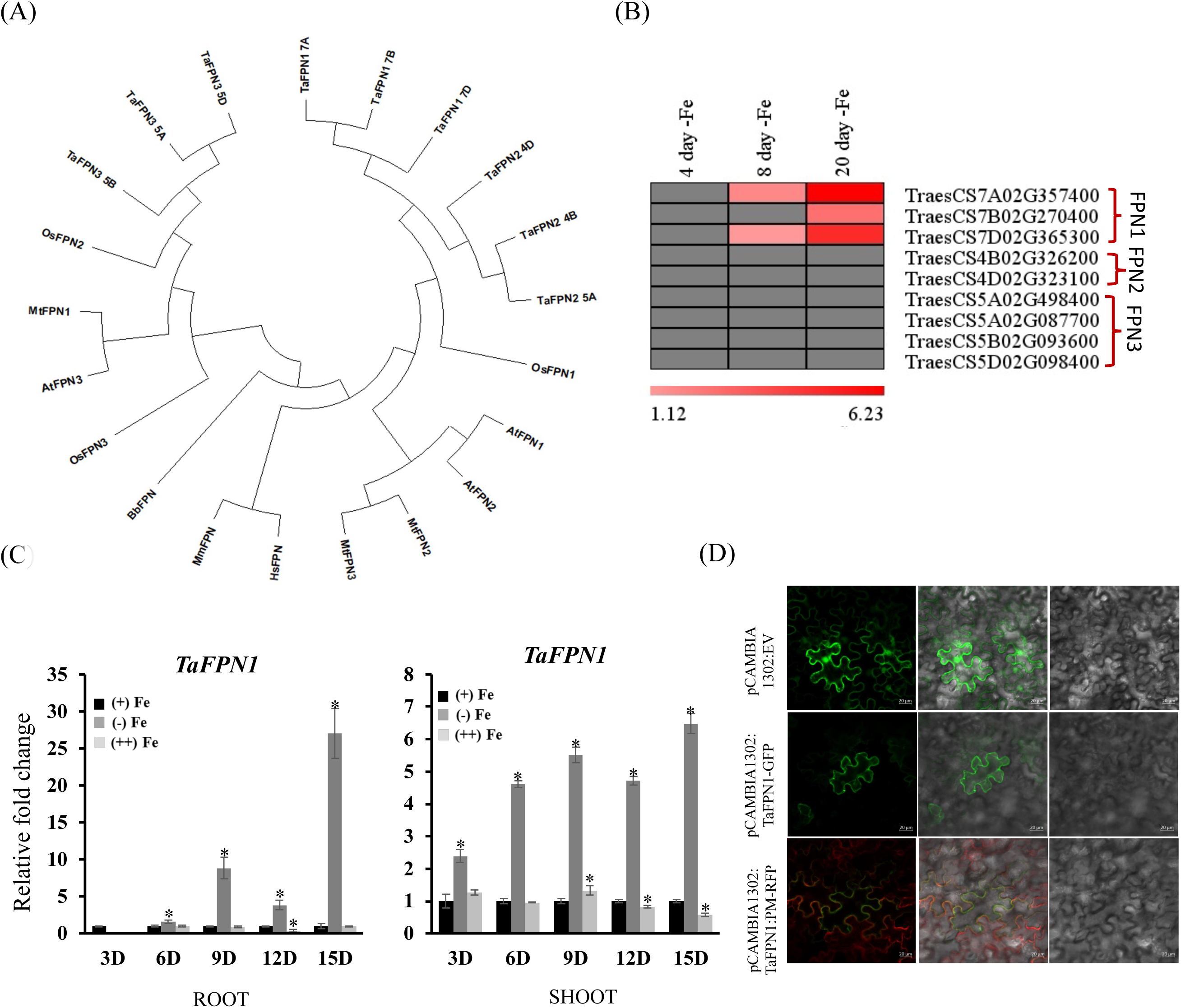
Phylogenetic analysis and characterization of the wheat Ferroportins. (A) Phylogenetic relationship between the identified wheat FPN proteins along with FPN proteins from *Arabidopsis thaliana*; AT2G38460 (AtFPN1), AT5G03570 (AtFPN2), AT5G26820 (AtFPN3), *Oryza sativa*; Os06t0560000 (OsFPN1), Os12t0562100 (OsFPN2), Os05g0131500 (OsFPN3), *Medicago truncatula*; Medtr1g084140 (MtFPN1), Medtr4g013240 (MtFPN2), Medtr4g013245 (MtFPN3), *Homo sapiens* (NP_055400.1), *Mus musculus* (NP_058613.2) and *Bdellovibrio bacteriovorus* (WP_011164469.1). The protein sequences were aligned using MUSCLE algorithm and unrooted phylogenetic tree was constructed via neighbour-joining method using MEGA 11. Bootstrap values were calculated as a percentage of 1000 trial. (B) Expression profile/s of wheat *FPN* genes resulted via transcriptome analysis of wheat roots under 4, 8 and 20 Days of Fe deficiency. The log_2_FC values of the respective genes were used to generate the heatmap using MeV software (version 4.9). (C) Expression analysis of *TaFPN1* in roots and shoots of wheat seedlings subjected to Fe deficiency and Fe surplus treatments for 3D, 6D, 9D, 12D, and 15D. C_t_ values were normalized against wheat *TaARF1* as an internal control. Data represent the mean of three biological replicates. Vertical bars represent the standard deviation. * on the bar indicates that the mean is significantly different at p < 0.05 with respect to their respective control (n=3). (D) Transient expression of *TaFPN1-GFP* in *N. benthamiana* leaf epidermal cells. Images were acquired at 48 h post agroinfiltration using LEICA DM600CS confocal microscope and processed with Fiji-ImageJ. GFP fluorescence was detected at 500–530 nm, whereas the red fluorescence was detected at 580–630 nm. Top panel = pCAMBIA1302 empty vector, Middle panel = *TaFPN1-GFP*, Bottom panel = Overlay of *TaFPN1-GFP* with plasma membrane marker : pMRK-RFP. Scale barL=L20 μm.

### Spatio-temporal expression characterization of TaFPN1

To have an insight into the possible function of *TaFPN1* in long-distance transport, a tissue-specific expression study was done in roots, stem, leaf, and flag leaf at 14 DAA. qRT-PCR . expression analysis suggested high expression of *TaFPN1* s in roots, and the detection of the transcript was also observed in the stem, leaf, and at a very low level in the flag leaf (Figure 2A). This suggests that the root may be the primary site of the *TaFPN1* function. Our *in-silico* expression survey also supports the root-specific expression (Figure S2A). Further, expression was also studied in the developing wheat grains, and a high gene expression was observed with the grain maturation (Figure 2B). *TaFPN1* expression was also found in the aleurone layer of the wheat grains (Figure S2B & Figure 2C). To address the cell-type-specific function of the wheat *FPN1*, we utilized the dataset generated from the wheat root tip (Zhang et al. 2023). Our analysis indicates the expression of the *TaFPN1* in the endodermis, suggesting its role in the radial transport of Fe. Our wheat exVip expression analysis suggested high expression of *TaFPN1-7A* among all the homoeologs (Figure S2C).

**Figure 2:**
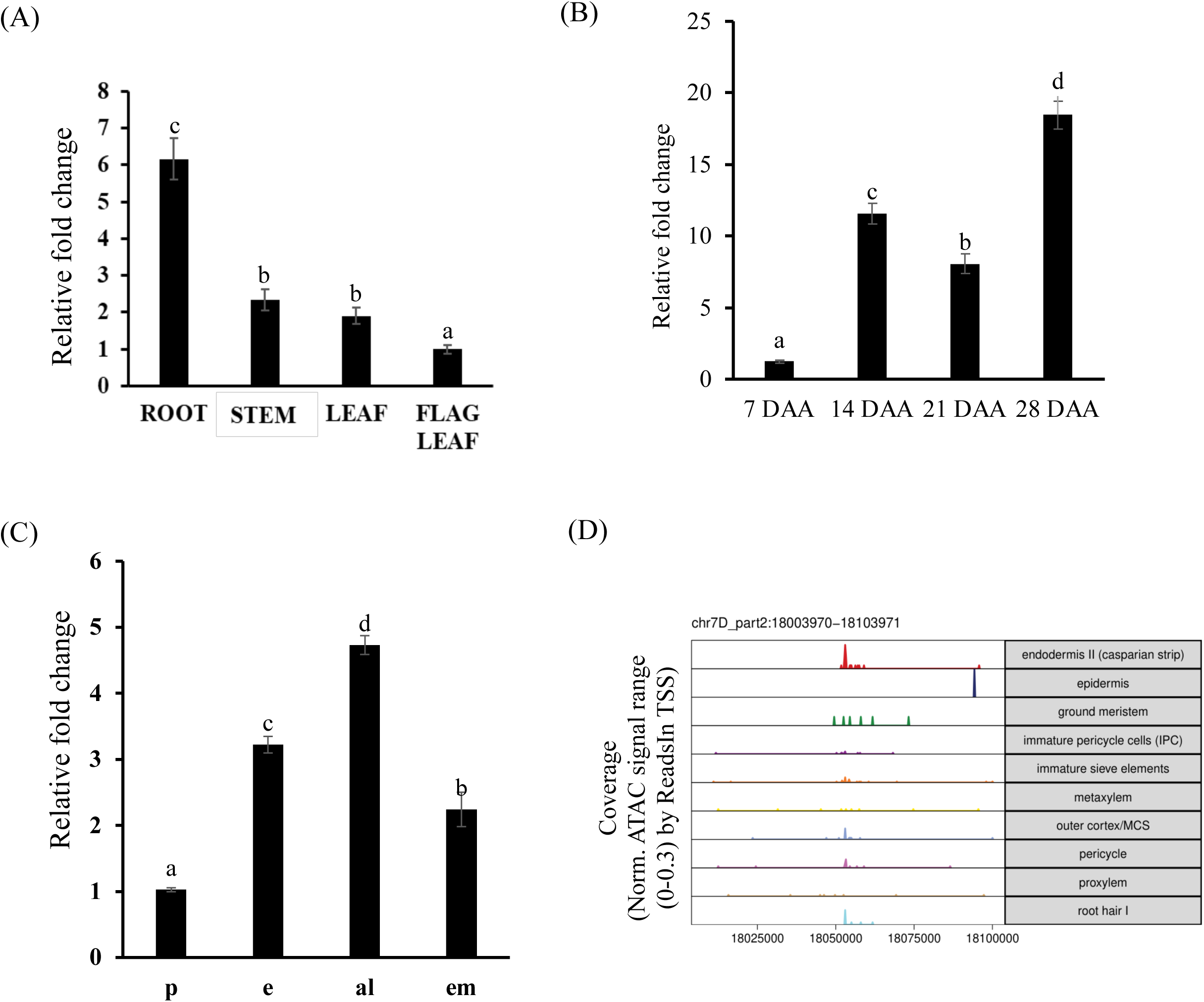
Expression characterization of *TaFPN1* in different wheat tissues. (A) Expression of *TaFPN1* in different wheat tissue harvested at 14 days after anthesis plants. (B) Exprerssion analysis of *TaFPN1* during wheat grains development during mentioned days after anthesis (DAA). (C) Expression analysis of *TaFPN1* in different tissue of grains (14 DAA stage). p- pericarp; al-aleurone; em-embryo; e-endosperm. The expression was normalized with *TaARF1* as internal control and fold change was calculated via 2^−ΔΔCT^ method. Vertical bars represent the standard deviation (n=3). Letters above the bars indicate significant difference (P<0.05) between the different tissues by one way ANOVA test. (D) Cell-type analysis of *TaFPN1-7D* (representative homeolog) using the normalized ATAC signal reads as obtained by using wheat single cell-type expression atlas in different root tissue (Zhang et al. 2023).

### TaFPN1 complements the sensitivity of atfpn1 for Fe deficiency and Fe surplus

To determine the role of *TaFPN1* in Fe homeostasis, we overexpressed *TaFPN1* in the *Arabidopsis* knockout mutant, *fpn1-2* (SALK_145458) - a sequence-indexed T-DNA insertion line. This was achieved by transforming homozygous *fpn1-2* (SALK_145458) plants using the *TaFPN1*-pGWB520 plant overexpression construct. Homozygous T3 lines showing 100% resistance to hygromycin B were screened and subjected to qRT-PCR analysis of *TaFPN1*. As a result, two transgenic lines (Line#2 and Line#8) displayed relatively higher expression and therefore, were selected for further characterization (Figure S3A).

Because FPN1 is known to influence Fe transit from root to shoot, *TaFPN1* was investigated for long-distance Fe transport in transgenic plants. To check whether iron was reaching the shoot efficiently, shoot chlorophyll concentrations were measured. Transgenic *Arabidopsis* plants, including Col-0 and *fpn1-2* mutant were grown on B5 medium for 5 days at 22 ± 1°C, 16 h light/8 h dark cycle, and 100μmol photons m^-2^s^-1^. They were then transferred to minimal media that was either Fe sufficient (FeIII-EDTA 50μM), Fe deficient (0 μM FeIII-EDTA + 300μM ferrozine), or Fe surplus (FeIII-EDTA 200μM). The shoots were harvested and tested for chlorophyll content, as described in the experimental procedures. It was observed that the *atfpn1* mutant had impeded iron mobility and was chlorotic, having decreased chlorophyll levels under all treatments (Figure 3A). However, *TaFPN1* overexpressing lines caused a reversal of the chlorotic phenotype in the mutant and resulted in a markedly significant increase in the chlorophyll contents under +Fe, –Fe, and ++Fe treatments (Figure 3B). Moreover, significant differences were observed in the primary root lengths of *TaFPN1* overexpressing lines compared to the *fpn1-2* mutant. The mutant had stunted root growth under all Fe conditions, whereas the *TaFPN1* overexpressing lines showed a significant increase in root lengths along with protruding lateral roots similar to Col-0 plants (Figure 3C). These findings suggest that *TaFPN1*-mediated Fe stress tolerance is associated with Fe mobilization from root to shoot.

**Figure 3:**
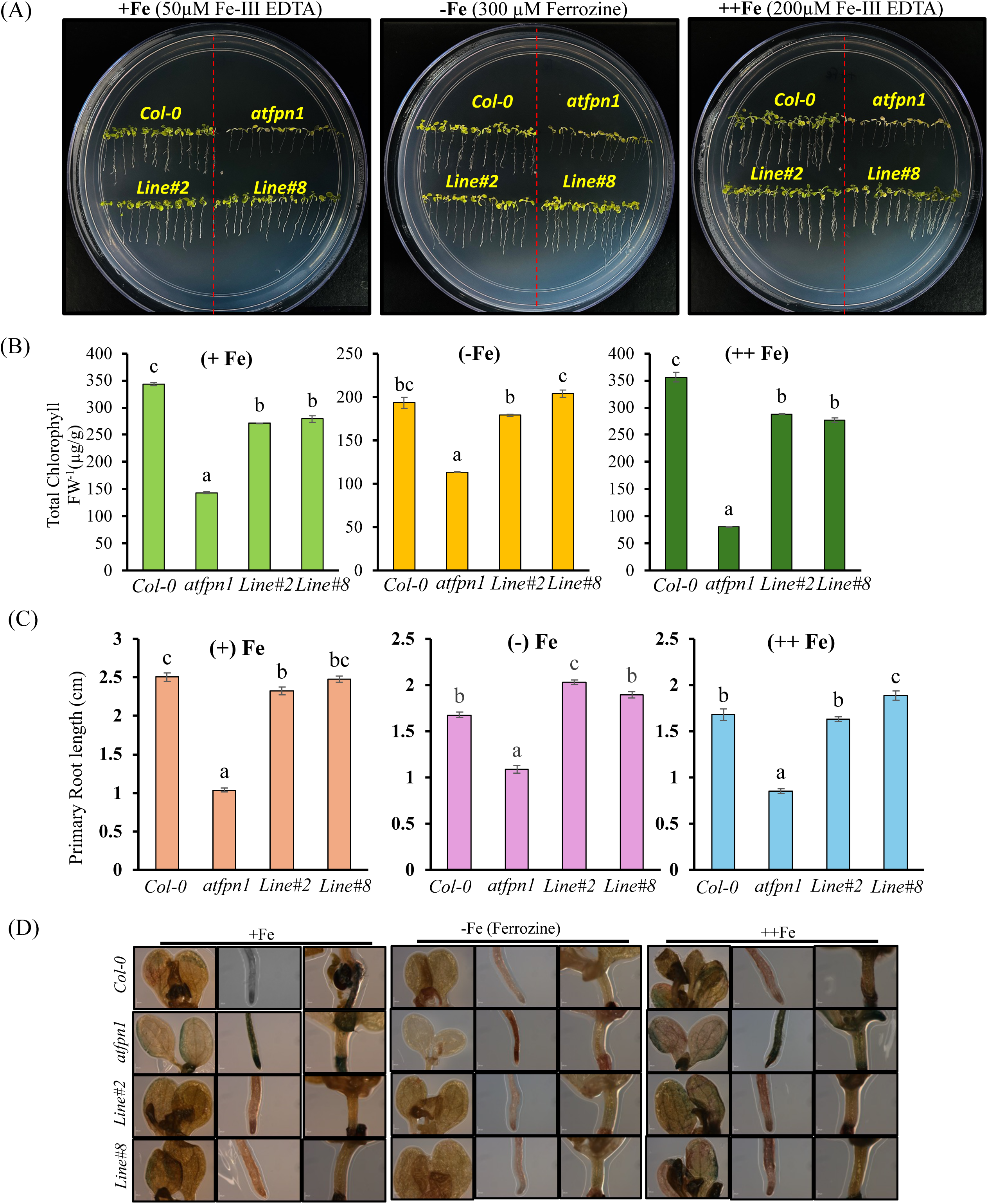
Overexpression of *TaFPN1* rescues the chlorotic phenotype of Arabidopsis knockout mutant *fpn1-2*. (A) Plants were grown on B5 solid medium for 5 days and then transferred to +Fe, −Fe or ++Fe media for 2 weeks. Under all conditions, the *fpn1-2* mutant displayed stunted growth and chlorotic phenotype as compared to Col-0 plants due to the absence of functional shoot Fe loader – Ferroportin 1. (B) Estimation of the total chlorophyll content in the leaves of the treated plants. Chlorophyll assessment was performed using 100mg of leaf tissue. (C) Primary root growth under different condition. Root lengths were measured using ImageJ software (Version 1.38) Values in the presented graphs represent three independent biological replicates. The letters above the bars indicate a significant difference (P<0.05) between the genotypes by one-way ANOVA test. (D) Iron distribution analysis in *TaFPN1* overexpressing lines. Perls/DAB staining was done for root (maturation zone), leaves and hypocotyl of Col-0, *fpn1-2* and *TaFPN1* overexpressing lines. *Arabidopsis* seedlings were germinated on Gamborg’s B5 agar media for 5 days and then shifted to B5 minimal media amended with +Fe (control), −Fe (Fe-deficient) and ++Fe (Fe excess) for 10 days. Scale bar: Leaves (500µm, Magnification 5X), Root & Hypocotyl (200µm, Magnification 10X).

To further investigate differences in iron distribution as a consequence of overexpressing *TaFPN1* in the *atfpn1* mutant, the plants grown in +Fe, -Fe, and ++Fe media were subjected to iron staining using the Perls’ method, enhanced with 3,3′- diaminobenzidine. The roots, leaves and hypocotyl regions of the plants were visualized to determine the Fe distribution. We observed dense iron staining in the *atfpn1* mutant roots under all Fe conditions (Figure 3D). As a result, the leaves and hypocotyl of the mutant plants displayed lower Fe accumulation. By contrast, the roots of Col-0 and *TaFPN1* transgenic lines showed lower intensity of iron staining than the mutant and significantly high Fe pools in the leaves and hypocotyl regions (Figure 3D). Furthermore, our ICP-MS data indicates that *fpn1-2* seedlings accumulate very low Fe compared to Col-0. However, Fe content was significantly regained in the two transgenic lines (Line#2 and Line#8) (Figure S3B). These results proved that the *TaFPN1* was able to rescue the mutant phenotype by mobilizing Fe from root to shoot in the transgenic lines.

Earlier, FPN1 was shown to impart tolerance to heavy metals (Co and Ni) (Morrissey et al. 2009; Kan et al. 2022). Therefore, gene expression of *TaFPN1* was studied in roots and shoots of wheat seedlings subjected to different heavy metals (Ni, Co, and Cd). The presence of these heavy metals shows a remarkable effect on root growth (Figure S3C). Our expression data compared to control show that the transcript accumulation of *TaFPN1* was reduced or remained unchanged under Ni and Cd stress but was significantly high in Co stress in roots. This was similar to *FPN1* of *Arabidopsis thaliana*, which has been reported to load Co from root to shoot, and *OsFPN1,* which is involved in the root-to-shoot translocation of Co and Ni. In contrast, the shoot specific high expression was observed in the presence of Ni and Cd (Figure S3C). Overall, our data suggests that wheat FPN1 is functionally active and could complement the chlorotic *atfpn1* mutant Further, it would be interesting to study how the *TaFPN1* is regulated.

### *TaFPN1* is regulated without hepcidin binding and IRE/IRP module

How plant *FPN1* turnover is regulated at the transcript or the protein level is unaddressed. In humans, hepcidin regulates the FPN protein turnover by post posttranslational mechanism through internalization and degradation. The binding site for the hepcidin is reported to be an extracellular loop in which the residue Cysteine 326 is required (Sham et al. 2005; Drakesmith et al. 2005; Billesbølle et al. 2020; Figure S4A,B&C). Plant FPN1, including in wheat, lacks this particular residue; therefore, it is unlikely that this mechanism exists in plants (Figure 4A). A plant ortholog of hepcidin has not been found. Therefore, we hypothesized that the Fe equilibrium in plants could be regulated by the second possible reported mode of regulation (Mode 2; Figure 4B), which involves an IRE (iron-responsive element)/IRP1 (iron-regulatory protein-1) iron switch or via miRNA-mediated control (Mode3).

**Figure 4:**
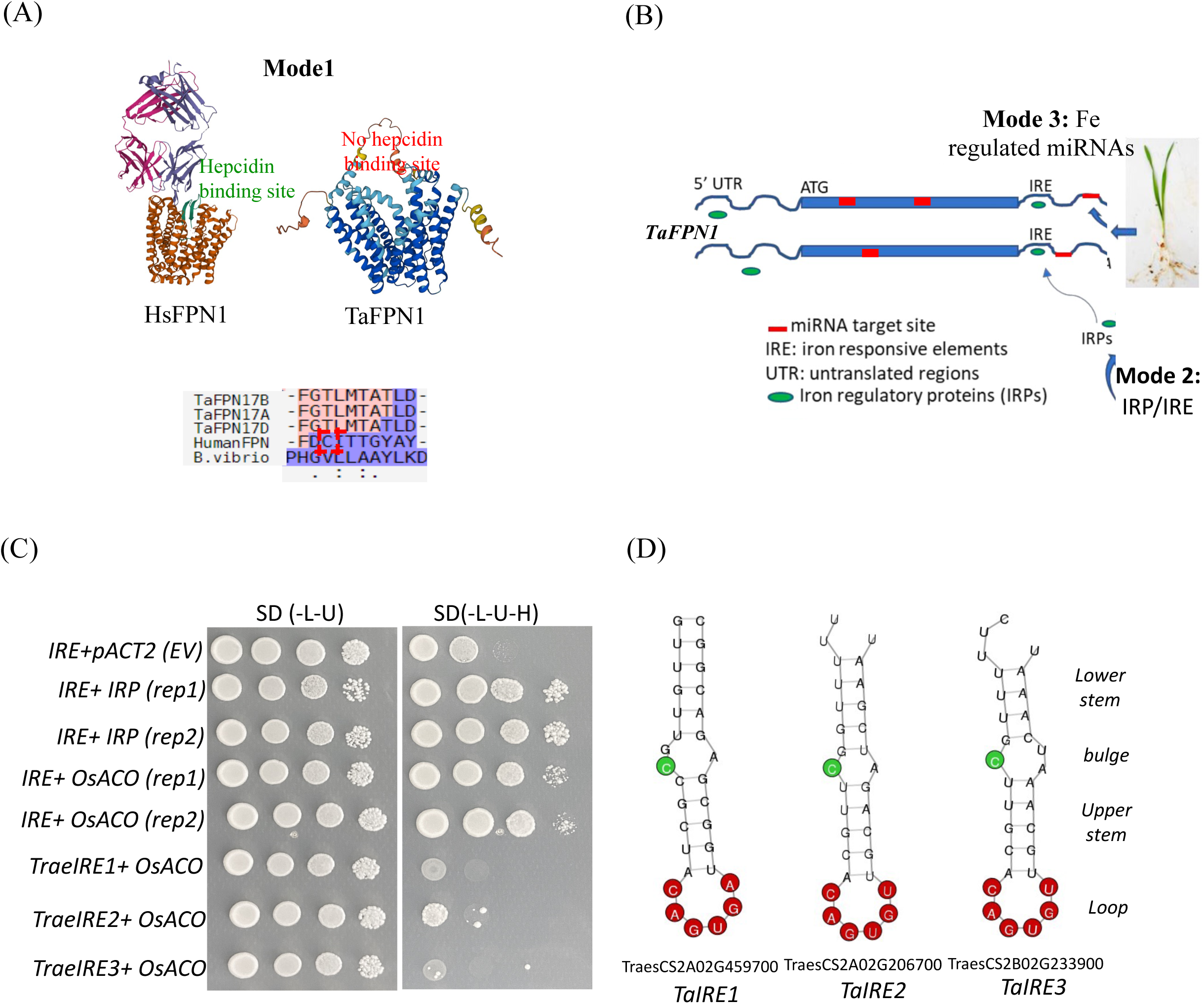
Regulation of *TaFPN1* at the post-transcriptional level: (A) The possible mechanism of regulation of *TaFPN1* could be either Mode1(hepcidin binding mediated). TaFPN1 lacks hepcidine binding site. HsFPN1 posses the hepcidin binding site as represented in the multiple sequence alignment. (B) Mode 2 (IRP-IRE mediated), or Mode3 (miRNA mediated). The *in-silico* structure of human and Wheat FPN1 reflects the presence of the hepcidin binding domain in HsFPN1 compared to TaFPN1. The lack of IRP/IRE elements in the 5’ and 3’ UTR regions of *TaFPN1* rules out this possibility of IRE-IRP mediated regulation. Multiple miRNA binding sites were observed for *TaFPN1*. (C) Yeast-three-hybrid interaction of IRE elements of human (HsIRE) and wheat (*TaIRE*) with human IRP (HsIRP) and rice Aconitase (OsACO). (D) IRE predicted the stability stem-loop structure for the different UTR of the wheat genes containing the conserved canonical structures. The bulge in all the and the line indicates the base pairing.

Recently, the rice Aconitase encoding gene *OsACO1* as a potential IRP protein was shown to be involved in the Fe regulatory mechanism in rice (Senoura et al. 2020). OsACO1 protein was speculated to have RNA binding ability, and because of this, the gene is hypothesized to be involved in post-transcriptional regulation of Fe homeostasis-related traits (Senoura et al. 2020). We hypothesized that OsACO1 participates in the RP/IRE-like module. Identifying the genes in wheat with potential IRE elements could identify multiple sites in the 5′ and 3′-UTR of Fe-regulated transcripts. Therefore we surveyed wheat genome promoters for the presence of the IRE (CxxxxxCAGUGNyyyyy) sites. Our *in-silico* analysis has revealed the presence of IREs in UTR regions of wheat genes, indicating the possibility of this mechanism in wheat (Table S2). Our searches for the presence of IRE elements in the 5 and 3′ UTR pf *TaFPN1* did not identify any IRE binding sites, ruling out the presence of UTR-mediated regulation.

Next, we checked if the IRE elements of wheat genes are regulated by human IRP or OsACO1. For this, we utilized the yeast three-hybrid reported system that could provide insight into such interactions. Previously reported human IRE/IRP was used as a positive control (Figure S5A). We selected three IRE sequences having high probability of interactions with low free energy scores (Figure S5B and Table S3). Our interaction assays revealed that OsACO1 could bind the human IRE sites (Figure 4C and Figure S5A). Multiple IRE elements showing high scores and stable stem-loop structures were cloned and checked for their interaction with OsACO1 and human IRP1 (Table S3). We then tested the binding of OsACO1 and human IRP with the wheat IRE sequences. We observed that OsACO1 does not show binding on the shortlisted wheat IRE sites. The promoter IRE element of wheat of different wheat transcripts shows a stable stem-loop structure (Figure 4D). Based on these data, we conclude that wheat FPN1 is not regulated by this mechanism predominant in humans (Mode 2). Therefore, we investigated the Mode 3 (miRNA-mediated) regulation of FPN1.

### Fe-induced MicroRNA miR1130 modulates gene expression of TaFPN1

MiRNA-mediated regulation has been demonstrated for multiple Fe-responsive genes, including human FPN. To investigate the miRNA-mediated mode of *TaFPN1* regulation, a small RNA sequencing dataset was generated from Fe-deficient root and shoot samples of wheat cv. “C-306” was employed (Sharma et al. 2023). Interestingly, among the microRNAs identified in the study, we were able to capture five miRNAs i.e., *tae-miR1130b-3p, tae-miR 1120c-5p, tae-miR1122c-3p, tae-miR1120a, tae-miR1120b-3p* targeting *TaFPN1* gene. To determine the strength of binding of these five miRNAs with *TaFPN1*, we scanned and performed searches for the putative miRNA binding site on the *TaFPN1* genomic DNA, cDNA, and CDS. Interestingly, *tae-miR1130b-3p* shows the best score (0) for the target prediction, suggesting that wheat FPN1 could possibly be controlled. *tae-miR1130b-3p* (Figure 5A). Therefore, we selected this miRNA to determine its antagonistic effect on *TaFPN1* gene. The vectors carrying the target site sequence,*TaFPN1:GUS* and the precursor sequence of *tae-miR1130b-3p*, were mobilized in tobacco leaves by agrobacterium mediated infiltration, and noted for reporter activity. A decline in the GUS activity was observed compared to the mock (Figure 5B&C). This decrease in the GUS activity in the presence of the miRNA validated *TaFPN1* as one of the targets and suggested that *tae-miR1130b-3p* negatively regulates *TaFPN1*. Most miRNAs in plant cells cleave target mature mRNAs; however, another regulatory mode, i.e., miRNA-intron interactions, has been predicted in *Arabidopsis* and *Oryza sativa* genes. These reports proposed the potential modes of action of plant miRNAs and gave conclusive proof for miRNA binding sites within the introns of the target genes. The miRNA binding sites in *TaFPN1* were also present in the intronic region.

**Figure 5:**
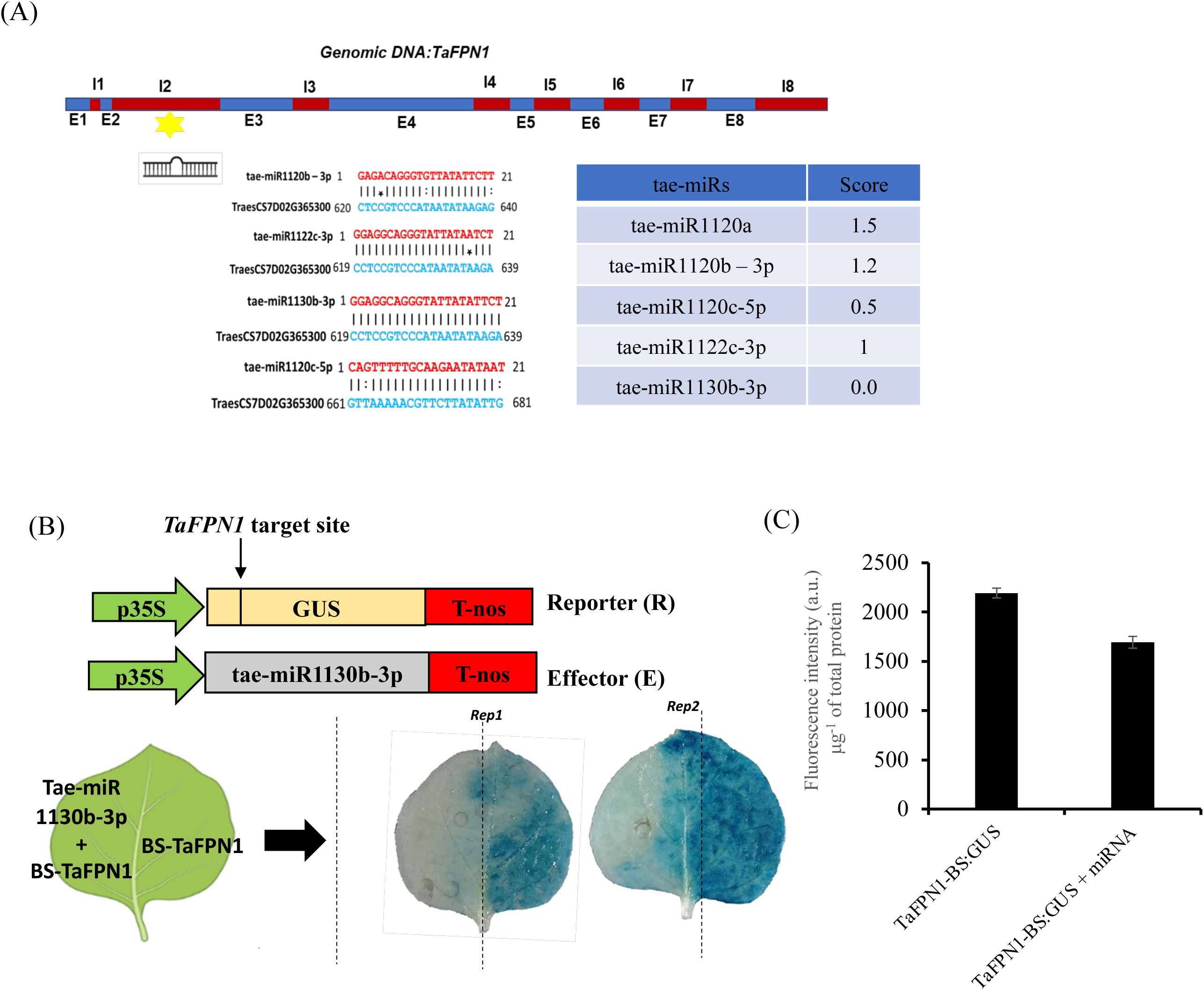
miRNA-mediated regulation of *TaFPN1* at the post-transcriptional level. (A) Exon-intron (Blue and red respectively) arrangement of the *TaFPN1* and the shortlisted candidate miRNA (score<1.5) capable of targeting *TaFPN1*. (B) Vector construction for co- transformation of target site and miRNA and in-vivo validation of the ability of *tae-miR1130b-3p* for *TaFPN1*. The intensity of the GUS (blue coloration) indicated the reporter’s expression. In contrast, the disappearance of the GUS indicates the degradation of the *TaFPN1:GUS* transcript, leading to a significant decrease in reporter activity. (C) GUS reporter activity as measured in 3 biological replicates (n=3 technical replicates).

### Spatio-temporal expression profiling of miRNA targeting TaFPN1

To understand the time-dependent regulation of the miRNAs targeting the *TaFPN1* gene, we studied their expression patterns in either of the tissues after subjecting them to Fe deficiency and Fe surplus stress for 3, 6, 9, 12, and 15 days. The analysis revealed that unlike the *TaFPN1* transcript, which accumulated under Fe deficiency and decreased under Fe excess compared to control Fe treatment, the opposite pattern was observed for the respective miRNAs under (– Fe) and (++) Fe conditions both in shoot and root. Overall, the transcripts of these miRNAs showed significant induction under Fe excess regimes, suggesting that molecular responses were established to control the accumulation of *TaFPN1* mRNA under Fe excess. Altogether, this approach helped us to characterize the tissue-specific expression responses into three categories. The first includes early responsive miRNAs, where *tae- miR1120c-5p* and *miR1122c-3p* showed early induction in the root, though, in the shoot, no miRNA showed early response (Figure 6A). The second category comprises late responsive *miRNA1130b-3p*, which mainly accumulated in root and shoot tissues. In shoot tissues, *tae- miR1120c-5p* also showed a late responsive nature in response to Fe excess. The third category highlights the miRNAs with mixed temporal expression during Fe excess, which included *tae-miR1122c-3p* in shoot and *tae-miR1120b-3p* in root and *tae-miR1120a* in both root and shoot (Figure 6A&B). We observed a high degree of inverse correlation between the miRNA and the *TaFPN1* expression under specific time points of Fe stress treatment. For example, in roots the miRNAs were found to be negatively correlated with *TaFPN1* expression at all the Fe stress time points but the highest correlation was observed for *tae- miR1130b-3p* at 6D with an R^2^= -0.97 (Figure 6C). Similarly, other miRNAs in root tissue also displayed significant negative correlation in the order *tae-miR1122c-3p* (R^2^= -0.92 at 12D) > *tae-miR1120c-5p* (R^2^= -0.72 at 9D) > *tae-miR1120b-3p* (R^2^= -0.67 at 12D) and *tae- miR1120a* (R^2^= -0.63 at 12D). In shoots, *tae-miR1120c-5p* displayed the highest R2 value of - 0.90 at 6D, followed by *tae-miR1120a* (R^2^= -0.62 at 9D), *tae-miR1130b-3p* (R^2^= -0.52 at 15 D) and lowest was observed for *tae-miR1122c-3p* (R^2^= -0.27 at 6D). This suggests that the expression of miRNAs is tissue and time-specific, wherein *tae-miR1130b-3p* in root and *tae- miR1120c-5p* in shoot could be the possible miRNAs in controlling the expression of *TaFPN1* gene, respectively. This explains that *miRNA1130p* can target *TaFPN1*, and the expression response represents these dynamic changes in the transcript during Fe stress. Our data suggests that *TaFPN1* is targeted by specific miRNA and is differentially induced during changing regimes of Fe in the rhizosphere.

**Figure 6:**
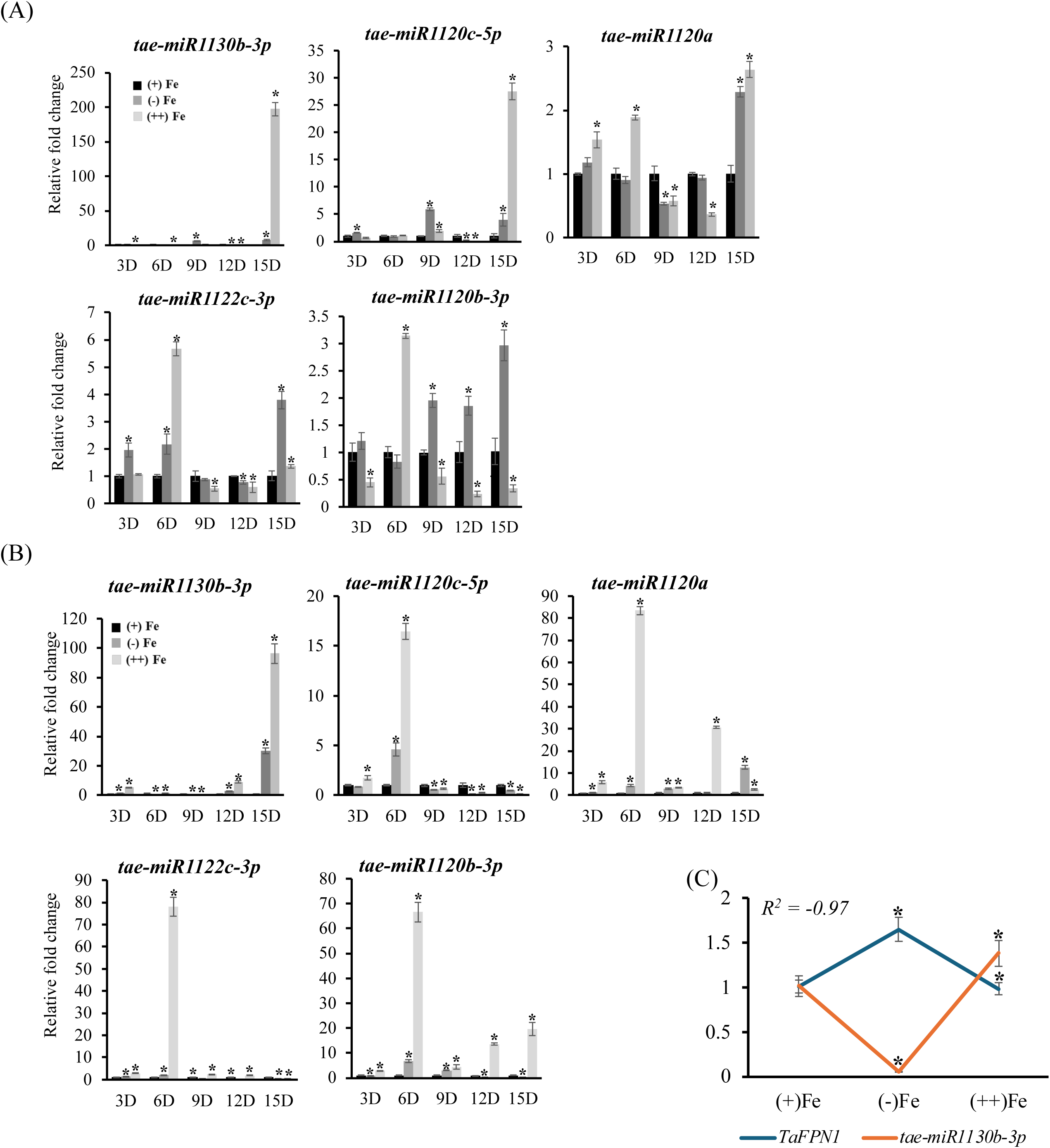
Temporal expression profiling of Fe-deficiency responsive miRNAs identified against *TaFPN1* in wheat. Expression profile of miRNAs in (A) root and (B) shoot tissues of wheat grown under Fe deficiency and Fe excess for different time points using qRT-PCR. Relative expression levels were calculated w.r.t. wheat reference gene *U6* snRNA transcript abundance (CT_gene_/CT_U6_) nL=L3; two-tailed Student’s t test (pL≤L0.05) was used to determine the significant change. (C) Correlation analysis between the expression of *TaFPN1* and *tae-miR1130b-3p* during Fe deficiency and surplus conditions in roots. Data represent the mean of three biological replicates. Vertical bars represent the standard deviation. * on the bar indicates that the mean is significantly different at p < 0.05 with respect to their respective control (n=3).

## Discussion

The current study investigates the functionality of plasmamembrane localized wheat Ferroportin 1 (*TaFPN1*) and the regulation of its turnover by *tae-miR1130b-3p* during iron homeostasis. Previously, the plasma membrane-localized FPN1 of *Arabidopsis thaliana and Oryza sativa* have been demonstrated to govern the root-to-shoot transit of Fe besides Co and Ni (Morrissey et al. 2009; Kan et al. 2022). However, the expression of *FPN1* in these plants was not iron-regulated and was found to be in low abundance in the roots of these plants. In contrast, analysis of our transcriptomic datasets of wheat roots at 4, 8, and 20 days of Fe deficiency revealed a significant accumulation of *TaFPN1* transcripts during the changing regimes of rhizospheric Fe (Kaur et al. 2019; Kaur et al. 2023). This was further corroborated by high temporal induction of *TaFPN1* transcripts in roots and shoots during Fe deficiency.

This suggests that *TaFPN1* might be helping to increase the shoot iron content by increasing iron loading into the root xylem and promoting iron efflux from the plasma membrane in the shoots. Expression of *TaFPN1* was found to be highest in wheat roots during the grain filling stage (14 DAA) and aligned with our analyses of *TaFPN1* for the tissue-specific expression using *in-silico* approaches. Additionally, the cell-type specific data showed high expression of *TaFPN1* in endodermis II (Casparian strips), which indicates its putative functionality as a metal ion effluxer (Zhang et al. 2023).

Our study observed that *TaFPN1* rescues the physiological symptoms of *fpn1-2* Salk mutant plants at different Fe concentrations. The chlorophyll content and the primary root lengths of *TaFPN1* expressing transgenic *Arabidopsis* plants showed high recovery comparable to wild-type Col-0 plants. Furthermore, differences in the iron distribution of mutant versus transgenic plants under +Fe, –Fe, and ++Fe revealed dense iron staining in the *atfpn1* mutant roots under all Fe conditions. and lower Fe accumulation in leaves and hypocotyl. However, Fe accumulation was low in the roots of Col-0 and *TaFPN1* transgenic plants but significantly high in the leaves and hypocotyl regions. Taken together, ICP-MS analysis revealed that the loss of function mutant *fpn1-2* displayed decreased Fe accumulation with respect to control. In contrast, *TaFPN1* overexpressing lines accumulated significantly higher Fe. This lends credence to the theory that FPN1 transports iron from the root, and its absence permits Fe to accumulate in the roots, thus providing Fe stress tolerance.

Our investigation of *TaFPN1* regulation revealed a miRNA-mediated control of its expression. The domain analysis of TaFPN1 with HsFPN1 at the amino acid level revealed conserved residues between both proteins. The Asp 39 and Asp 181 residues of HsFPN are known to be critical for Fe transport (Li et al. 2020). Asp181, which constitutes part of the Fe translocation was found to be highly preserved between the TaFPN1 and HsFPN protein sequences (Figure S1). However, Ala was present in lieu of Asp39 in TaFPN1, suggesting that this residue might not be essential for iron transport by TaFPN1. In HsFPN, Asn174, Asp 325, and Arg466, Tyr 318, Tyr 331, and Tyr 501 are also known to affect iron efflux, although not directly as a Fe-binding site (Taniguchi et al. 2015; Bonaccorsi di Patti et al. 2015; Billesbølle et al. 2020). Among these, Tyr 318, Asn174, and Arg466 were highly conserved, whereas Tyr 501 displayed synonymous amino acid substitution of Ser in TaFPN1 protein sequences. Similarly, Tyr 331 was also replaced with its synonymous amino acid Thr in TaFPN1. Overall, the conservation of several residues that are known to be directly or indirectly involved in Fe binding/transport in wheat FPN1 suggests that TaFPN1 is likely to transport of Fe^2+^, which is a primary substrate of FPN orthologs (Drakesmith et al. 2015; Li et al. 2020).

Despite significant conservation among the Fe binding and transport residues between wheat and human FPN orthologs, the absence of Hepcidin binding residue in wheat FPN1, including wheat and *Arabidopsis,* confirms the absence of this regulatory mechanism. Therefore, miRNA-mediated regulation of the *TaFPN1* was explored by identifying the miRNA binding sites. Earlier, human miR-485-3p was shown to bind 3’ UTR of human *FPN1* at post-transcriptional level. Our previous small RNA dataset of Fe deficiency exposed wheat roots and identified multiple miRNAs that could target FPN1 (Sharma et al. 2023).

Spatio-temporal stem-loop qRT-PCR expression analysis of these miRNAs under Fe deficiency and surplus showed significant induction under Fe excess regimes. We observed a high inverse correlation between the miRNA and the TaFPN1 transcript expression during Fe stress treatments. In roots, the highest correlation was observed for *tae-miR1130b-3p* with an R^2^= -0.97. Furthermore, *tae-miR1130b-3p* was found to negatively regulate *TaFPN1* gene when co-expressed with *TaFPN1:GUS* in tobacco leaves. During Fe stress treatments, we observed a high inverse correlation between the miRNA and the TaFPN1 expression*. Tae- miR1130b-3p* shows a binding site in the intronic region of the *TaFPN1*. It could be possible that the region is part of the pre-mRNA. Using bioinformatic approaches coupled with degradome analysis, a large-scale miRNA binding was present in the introns of *Arabidopsis* and rice (Meng et al. 2013). miRNA-labelled assays point to identifying its targets located not only in the promoter but also in an intron, exon, and intergenic (Xun et al. 2020). Although the mechanism for the intronic targets is not clear, one could speculate the role of *tae-miR1130b-3p* in generating splice variants. Whether *TaFPN1* forms different splice variants and its role needs to be investigated. A decline in the GUS activity was observed, suggesting that *tae-miR1130b-3p* specifically targets *FPN1* for its degradation.

Human *FPN* mRNA contains an iron-responsive element (IRE) motif; likewise, human DMT1, ferritin, and the transferrin receptor, FPN is subject to Fe-dependent regulation by interacting iron regulatory proteins (IRPs) with the IRE sequence (Hentze et al. 2010; Kühn 2015). Identifying the genes in wheat with potential IRE elements revealed multiple sites in the 5′ and 3′-UTR of Fe-regulated transcripts. Interestingly, none of the wheat IRE sites interacted with the human IRP or OsACO1. We conclude that it is possible that such an IRE/IRP mechanism does not exist in plants. It has been reported that rice mutant for *ACO1* showed delayed Fe dependent molecular response. At this point, it would be interesting to explore whether these Fe homeostasis genes show the presence of IRE elements in their promoters. Interestingly, no changes in the expression of the iron deficiency-responsive cis-acting element binding factors 1 and 2 (IDEF1/2) were seen in rice plants (Senoura et al. 2020). In contrast, *Arabidopsis ACO1* mutants showed no molecular changes for Fe homeostasis genes, including ferritin (Arnaud et al. 2007). Therefore, the role of ACO1 as a potential IRP still needs investigation. Alternatively, the delayed response of Fe homeostasis genes in *osaco1* mutant could be via other pathways, yet further experimentation is needed. It is also unclear about the RNA binding activity of plant ACO, which could result from either the structural element or the presence of the canonical sequence. Based on current evidence, we conclude that miRNA-mediated regulation of the *FPN1* transcript is well conserved among plant and animal kingdoms. Overall, this study unravels the physiological role of wheat FPN1 and provides functional evidence of its participation in Fe homeostasis along with miRNA-mediated regulatory control of its expression.

## Acknowledgments

The authors thank the Executive Director of NABI for the facilities and support. This work was funded by the Department of Science and Technology- Science & Engineering Research Board (SERB) Grant number CRG/2020/000940 and the NABI-CORE grant to AKP. DBT-eLibrary Consortium (DeLCON) is acknowledged for providing timely support and access to e-resources for this work.

## Author Contribution

AKP and SS conceived and designed the research. SS performed most of the wet lab experiments and data analyses. SS, RJ and APS conducted other wet lab experiments; AKP and SS wrote and finalized the manuscript. All authors have read and approved the final manuscript.

## Declarations

### Ethics Approval and Consent to Participate

Not applicable

### Consent for Publication

All authors approved the manuscript for publication.

