## Supplementary Figures for "MicroRNA Tae-miR1130p targets wheat ferroportin1 (*TaFPN1*) in the absence of iron-responsive element/iron-regulatory protein1 module"

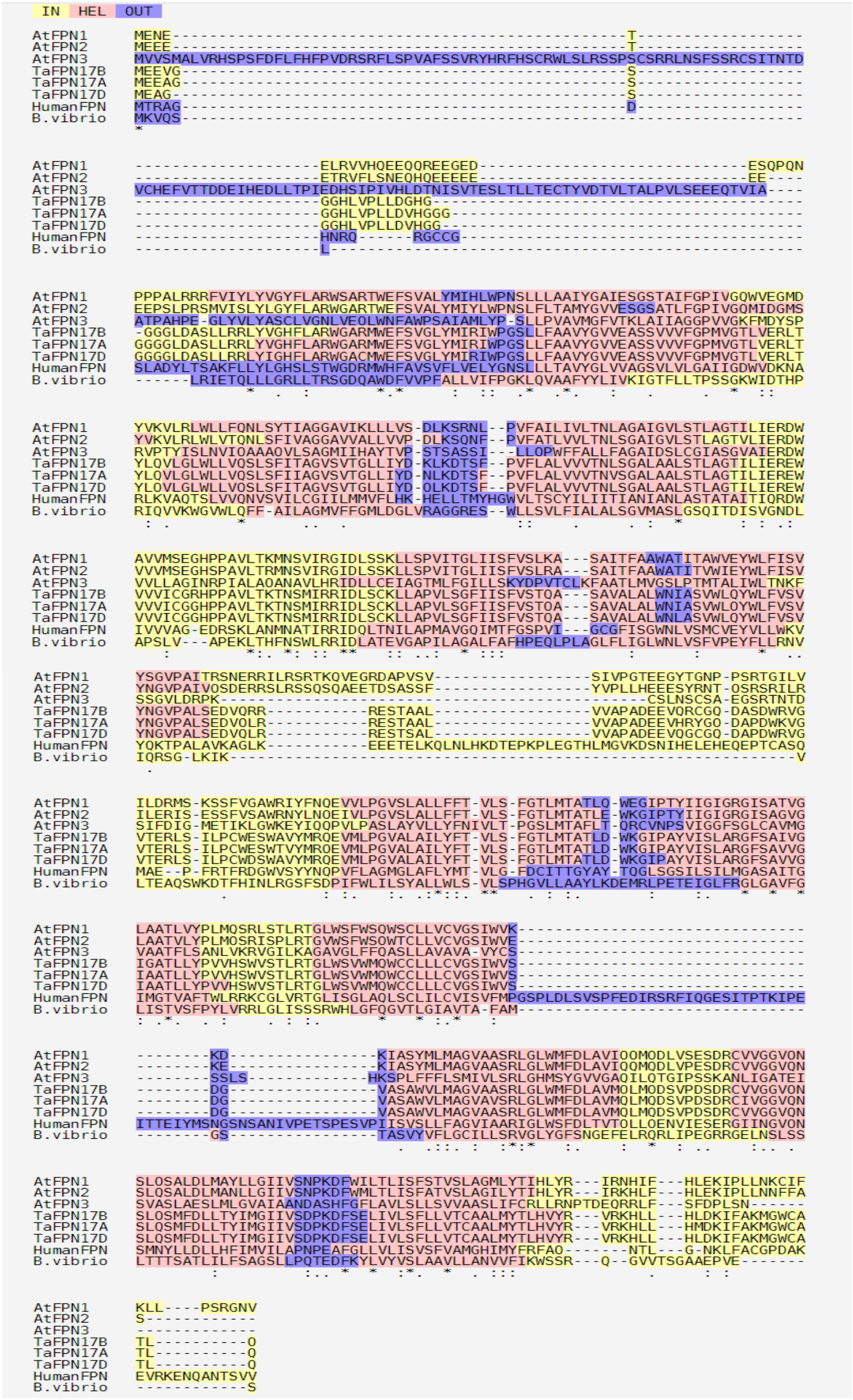


**Figure S1: Sequence alignment of different FPN1 proteins.** Protein sequence alignment between the identified *Triticum aestivum* FPN protein sequences and FPN proteins belonging to *Arabidopsis thaliana* (AtFPN1, AtFPN2, AtFPN3), *Homo sapiens* (HsFPN), and *Bdellovibrio bacteriovorus* (BbFPN). Multiple sequence alignment was done using T-Coffee (Di Tommaso et al. 2011). The shaded blocks in red represent sequence regions with strong alignment, while yellow blocks denote regions with moderate alignment. Blue-green blocks indicate regions with weak alignment.

(A)

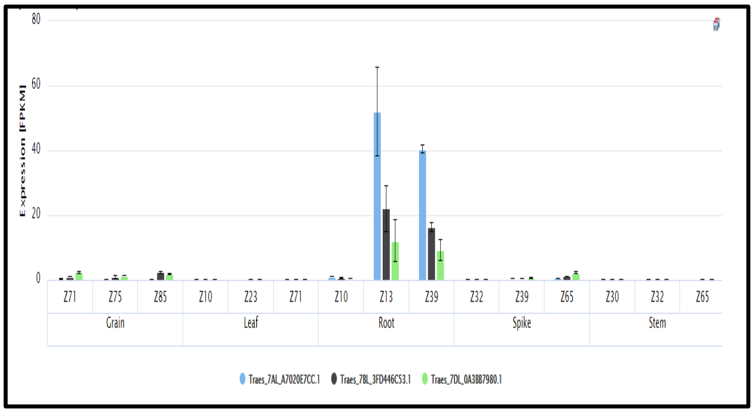


(B)

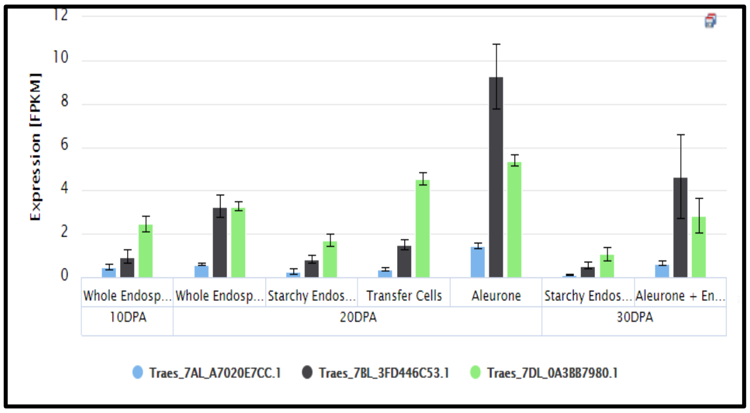


(C)

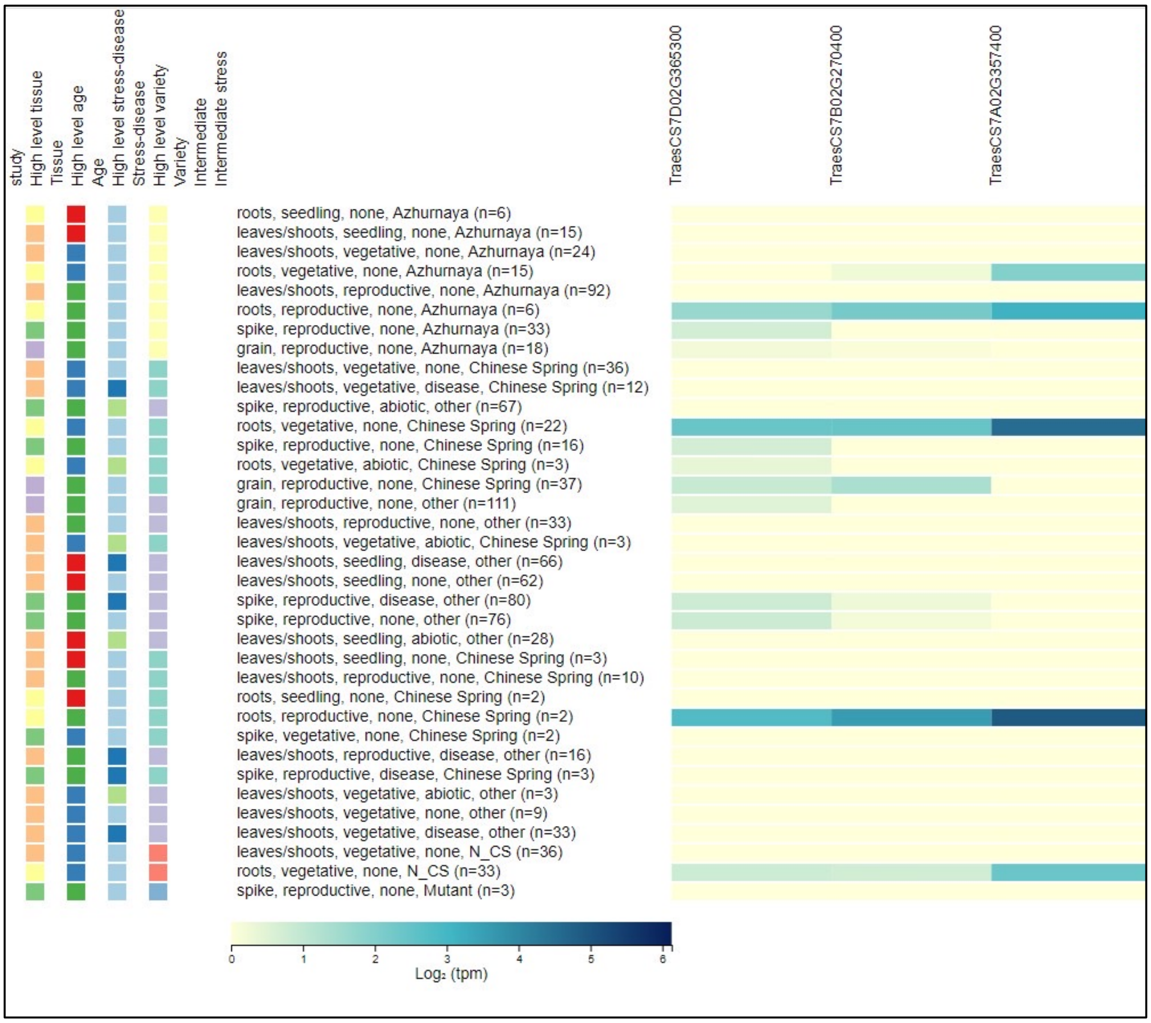


Figure S2: ***In silico* expression analysis of wheat FPN1 along with their homeologues:** (A) Different wheat tissues (spike, root, leaf, grain and stem) each sampled at three different developmental stages providing broad overview of gene expression during the developmental time course, the x-axis depicts the Zadoks growth scale where highest expression of TaFPN1 was seen as Z13(Three leaves visible) and Z39(Flag leaf ligule just visible). (B) Depicts TaFPN1 gene expression at Grain layers at various days post anthesis, where it shows that TaFPN1 is highly expressed at Aleurone 20 day post anthesis . (C) Gene expression analysis was performed using wheat expression database browser (http://www.wheat-expression.com/ (Wheat Exp)).

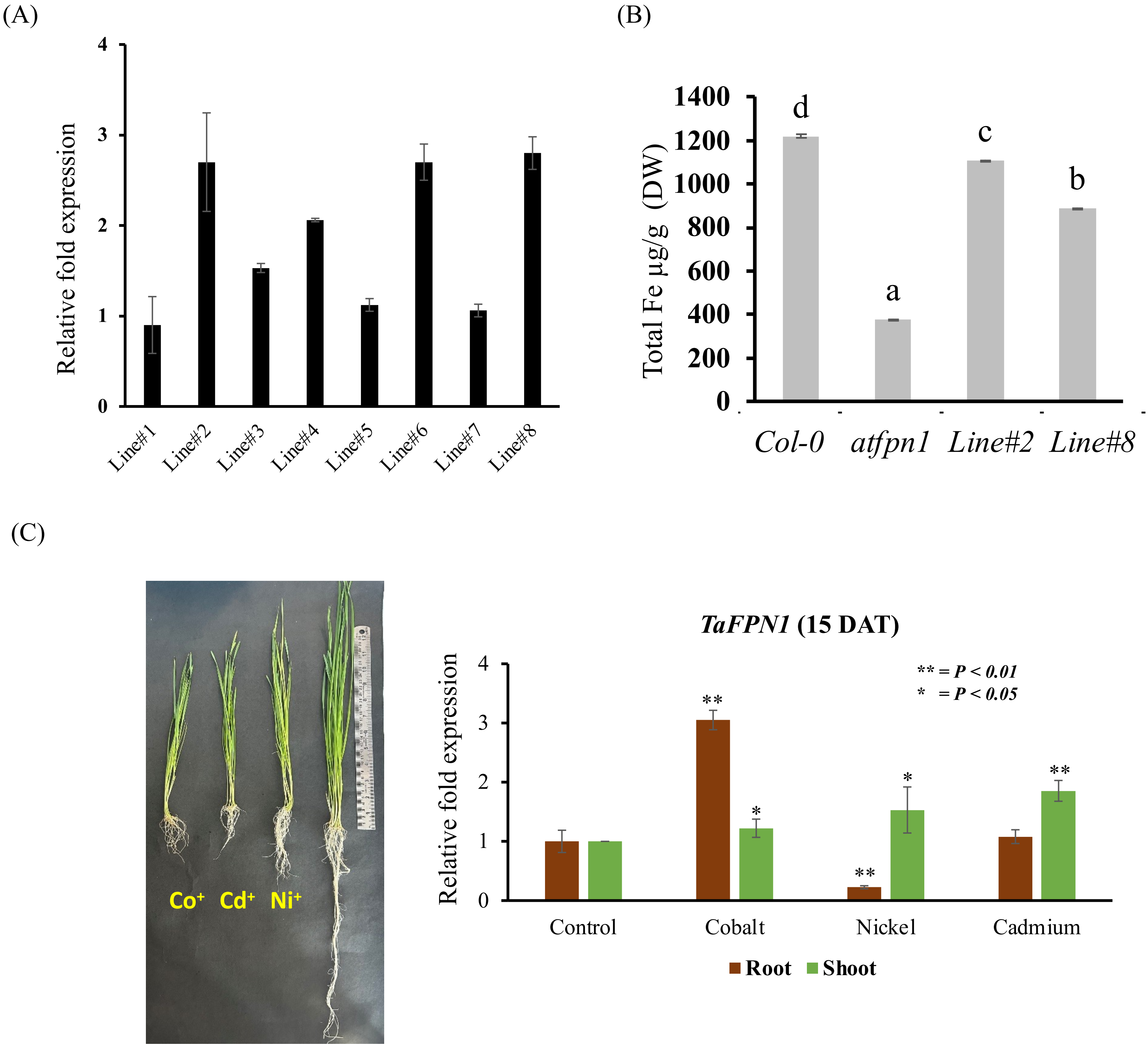


**Figure S3: Characterization of TaFPN1 complemented lines in *atfpn1* and its expression analysis under heavy metal stress.** (A) qRT-PCR expression analysis of *TaFPN1* transcript in *Arabidopsis* transgenic lines. 10-12 plants from each line were taken followed by RNA extraction and cDNA preparation. Data represent the mean of three biological replicates. Vertical bars represent the standard deviation. Letters indicate significant difference (P<0.05) between transgenic lines by the one way ANOVA test. Line#2 and Line#8 showed highest overexpression of the gene. (B) Fe accumulation changes in Col-0, *fpn1-2* and *TaFPN1* overexpressing lines analysed by ICP-MS under control conditions. The Data shown is an average of 3 biological replicates (n=3 technical replicates). Significant differences (P<0.05) were calculated by one-way ANOVA test and are represented by different letters above the bars. (C) Phenotypic data shows Effect of heavy metal stress on wheat shoot and roots. The plants showed toxicity symptoms in the form of stunted growth. Pictures of plants were taken after 15 days of treatment. (C) Phenotype of wheat seedlings subjected to heavy metal stress (right panel). Expression of *TaFPN1* genes in roots and shoots of wheat plants subjected to heavy metals stress for 15 days (left panel). The expression was normalized with *TaARF1* as internal control and fold change was calculated via $2^{-\Delta\Delta CT}$ method. Vertical bars represent the standard deviation; * represents the significant difference at p-value < 0.05 with respect to their respective control treatments (n=3).

(A)

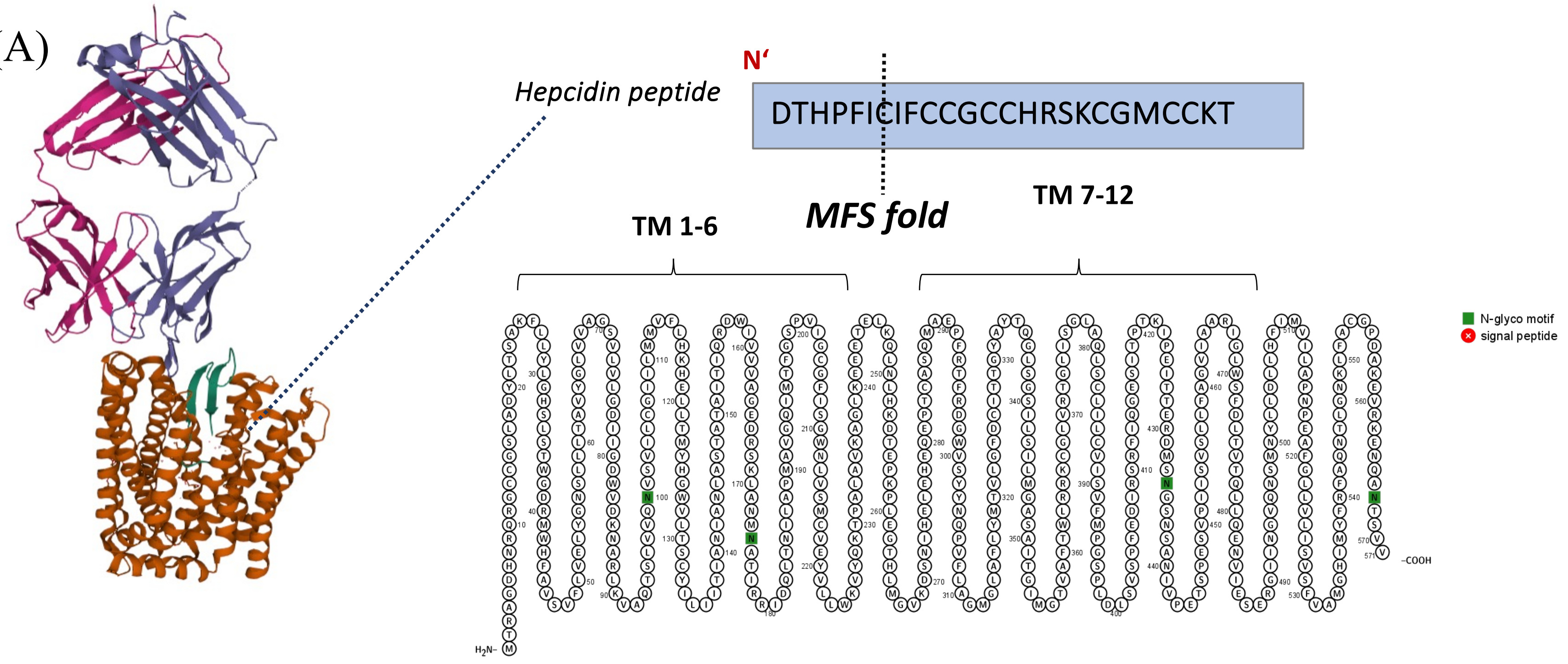


(B)

**Human FPN (571 amino acids)**

MTRAGDHNRQRGCCGSLADYLTSAKFLLYLGHSLSTWGDRMWHFAVSVFLVELYGNS

LLLTAVYGLVVAGSVLVLGDIIGDWVDKNARLKVAQTSLVVQNVSVILCGIILMMVFLHKHE

LLTMYHGWVLTSCYILIITIANIALASTATAITIQRDWIVVVAGEDRSKLANMNATIRRID

QLTNILAPMAVGQIMTFGSPVIGCGFISGWNLVSMCVEYVLLWKVYQKTPALAVKAGLKE

EETELKQLNLHKDTEPKPLEGTHLMGVKDSNIHELEHEQEPTCASQMAEPFRTFRDGWVS

YYNQPVFLAGMGLAFLYMTVLGFDCITTGYAYTQGLSGSILSILMGASAITGIMGTVAF

TWLRRKCGLVRTGLISGLAQLSCLILCVISVFMPGSPLDLSVSPFEDIRSRFIQGESITPTKIPEI

TTERDMSNGSNSANIVPETSPESVPIISVSLLFAGIVIAARGLWSFDLTVTQLLQENVIESER

GIINGVQNSMNYLLDLLHFIMVLAPNPEAFGLLVLISVSFVAMGHIMYFRFAQNTLGN

KLFACGPDAKEVRKENQANTSVV

**Figure S4: Protein domain analysis of human FPN1.** (A) Structure analysis of human FPN for the hepcidin binding site. The presence of 12 transmembrane domains is also reflected in the image. Human FPN bound to hepcidin (retrieved from PDB) b. Transmembrane structure of human FPN predicted by PROTTER (B) Amino acid sequence of human FPN-conserved Histidine 32 residue, Fe binding residues, Hepcidin binding site (cysteine 326), residues that decrease hepcidin binding if mutated, the enclosed boxes indicate transmembrane helices from 1 to 12.

(A)

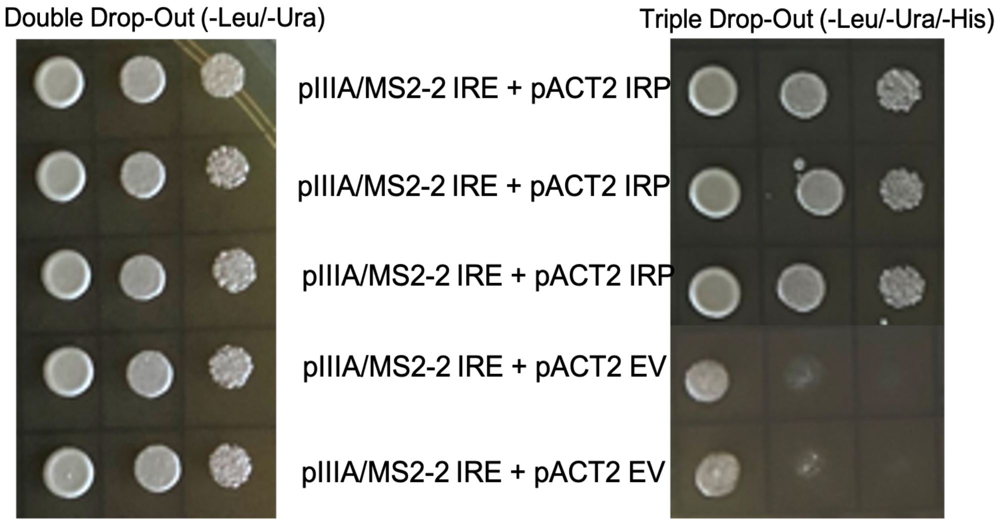


(B)

| Wheat gene ID | Predicted annotation | Free Energy Score | Quality |
|---|---|---|---|
| TraesCS2A02G206700 (3'UTR) | transmembrane protein | -9.90 | High |
| TraesCS2B02G233900 (3'UTR) | transmembrane protein | -7.90 | High |
| TraesCS6D02G402800 (5'UTR) | Zinc finger CCCH domain-containing protein 19 | -5.70 | High |
| TraesCS2B02G186800 (3'UTR) | Mitochondrial import inner membrane translocase subunit TIM44 | -3.70 | High |

(C)

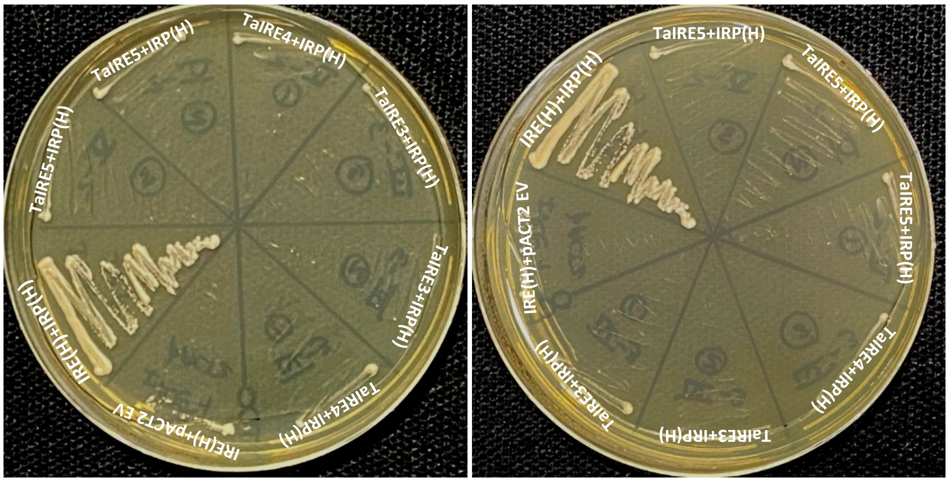


**Figure S5: Interaction studies of IRE, IRP and Aconitase.** (A) Yeast 3 Hybrid data depicting interaction between human IRP and human IRE along with it's negative control. This yeast three-hybrid (Y3H) method was employed in this investigation to determine whether or not the IRE-IRP interaction exists in the hexaploid wheat genome. (B) Table depicting a list of selected wheat genes containing the iron-responsive elements (IRE) in either 5' or 3' UTR. CxxxxxCAGUGNyyyyy IRE sequence was looked for in all UTRs chosen wheat IREs having higher score by using SIREs (http://www.sires-webserver.eu/), Searching for Iron-Responsive Elements, the bioinformatic program for the prediction of IREs. (C) Human IRP were cloned in the pACT2-AD vector was used for interaction with wheat IREs, and human IRE those were cloned in the pIIIA/MS2-2 vector. After being co-transformed in the YBZ-1 yeast strain, the produced constructs were plated on SD/-Leu/-Ura/-His plates. Pictures were taken after 4 days of incubation at 30 $^{o}$C .
