## Supplementary Tables for "MicroRNA Tae-miR1130p targets wheat ferroportin1 (*TaFPN1*) in the absence of iron-responsive element/iron-regulatory protein1 module"

| Table S1: List of Primers employed for cloning in this study. | |
|---|---|
| Gene name | **Primer Sequence (5'-3')** |
| *TaFPN1* | **F :** TCCGGAGGATCGACCTGAGCTG<br>**R :** GCTGGACGTCCTCGCTGAGAG |
| *TaFPN2* | **F:** TCGCCGGAACCATACTGATCGAGAG<br>**R:** ACACGAAGAGCCAGTACACCAGCC |
| *TaFPN3* | **F:** CTCGGGGCTCCAATAGTTGGCAAAC<br>**R:** GCACAATGAACCAAGGCCTGAGAAGC |
| *TaARF1* | **F:** TGATAGGGAACGTGTTGTTGAGGC<br>**R:** AGCCAGTCAAGACCCTCGTACAAC |
| *AtActin* | **F:** ACCCGATGGGCAAGTCATCACG<br>**R:** CATGCTGCTTGGTGCAAGTGCTGTG |
| **Stem-Loop qRT Primers for wheat miRNAs** | |
| **miRNA** | **Primer Sequence (5'-3')** |
| *tae-miR1130b-3p* | **RT :**GTCGTATCCAGTGCAGGGTCCGAGGTATTCGCACTGGATACGACCCTCCG<br>**F:** GCGGCGGTCTTATATTATGGGA |
| *tae-miR1120c-5p* | **RT :**<br>GTCGTATCCAGTGCAGGGTCCGAGGTATTCGCACTGGATACGACGTCAAA<br>**F:** GCGGCGGTAATATAAGAACGTT |
| *tae-miR1122c-3p* | **RT:** : GTCGTATCCAGTGCAGGGTCCGAGGTATTCGCACTGGATACGACCCTCCG<br>**F :** GCGGCGGTCTAATATTATGGGA |
| *tae-miR1120b – 3p* | **RT :**<br>GTCGTATCCAGTGCAGGGTCCGAGGTATTCGCACTGGATACGACCTCTGT<br>**F :**<br>GCGGCGGTTCTTATATTGTGGG |
| *tae-miR1120a* | **RT:** GTCGTATCCAGTGCAGGGTCCGAGGTATTCGCACTGGATACGACCTCCGT<br>**F:** CGGCGGACATTCTTATATTATGAG |
| *U6 snRNA* | **F :** CTTCGGGGACATCCGATAAA<br>**R :** GACCATTTCTCGATTTGTGC |
| *Universal Reverse* | GTGCAGGGTCCGAGGT |

**Table 2: List of predicted wheat genes containing the iron responsive elements (IRE) in either 5' or 3' UTR. CxxxxxCAGUGNyyyyy IRE sequence was looked for in all UTRs (x and y complementary as AU/UA/GC/CG/UG/GU using an in-house generated Perl script.**

| Wheat Gene ID | Predicted Annotation | pFam-IDs | Free Energy Score | Quality |
|---|---|---|---|---|
| **3'UTR IRE Motif** | | | | |
| 1. TraesCS2A02G206700 | transmembrane protein | | -9.90 | High |
| 2. TraesCS2B02G233900 | transmembrane protein | | -7.90 | High |
| 3. TraesCS6B02G331100 | DNA polymerase III PolC-type | PF02811 | -7.70 | High |
| 4. TraesCS2D02G209200 | transmembrane protein | | -7.30 | High |
| 5. TraesCS6B02G042800 | F-box family protein | PF00646 | -6.20 | High |
| 6. TraesCS2A02G006800 | Disease resistance protein (NBS-LRR class) family | PF00931 | -5.90 | High |
| 7. TraesCS2A02G322200 | E3 UFM1-protein ligase 1 homolog | | -4.80 | High |
| 8. TraesCS2B02G186800 | Mitochondrial import inner membrane translocase subunit TIM44 | PF04280 | -3.70 | High |
| 9. TraesCS5D02G355900 | Plant regulator RWP-RK family protein, putative | PF02042; PF00564 | -2.90 | High |
| 10. TraesCS6D02G036100 | Mitochondrial transcription termination factor-like | PF02536 | -2.80 | High |
| 11. TraesCS5B02G114200 | Cyclin B1 | | -4.30 | Medium |
| 12. TraesCS4D02G121800 | Kinase family protein | PF07714 | -4.20 | Medium |
| 13. TraesCS2D02G473700 | Homeobox-leucine zipper family protein | PF00046; PF01852 | -3.50 | Medium |
| 14. TraesCSU02G016400 | Autophagy-related protein 22-1 | | -2.30 | Medium |
| 15. TraesCS7B02G329400 | T-box transcription factor, putative (DUF863) | | -2.20 | Medium |
| 16. TraesCS5B02G385900 | BTB/POZ domain containing protein, expressed | PF00917; PF00651 | -2.10 | Medium |
| 17. TraesCS5A02G349500 | Plant regulator RWP-RK family protein, putative | PF02042; PF00564 | -1.80 | Medium |
| 18. TraesCS7D02G526200 | Receptor-like protein kinase | PF00069 | -1.70 | Medium |
| 19. TraesCS4A02G174100 | Zinc finger protein, putative | PF13912 | -0.90 | Medium |
| 20. TraesCS3A02G489100 | cysteine-rich/transmembrane domain A-like protein | PF12734 | -1.90 | Low |
| 21. TraesCS1B02G352300 | WD40 repeat-like protein | PF00400; PF04003 | -0.70 | Low |
| 22. TraesCS2D02G481900 | Anthocyanidin reductase | PF01370 | -0.70 | Low |
| 23. TraesCS7A02G561900 | NBS-LRR disease resistance protein | PF00931 | -0.30 | Low |
| 24. TraesCS1B02G257200 | Leucine-rich repeat receptor-like protein kinase family protein | PF12819 | 0.0 | Low |
| 25. TraesCS1D02G245800 | Leucine-rich repeat receptor-like protein kinase | PF12819 | 0.0 | Low |
| 26. TraesCS4D02G292900 | Tubulin-tyrosine ligase family protein | PF03133 | 0.0 | Low |
| 27. TraesCS5A02G045700 | Protoporphyrinogen oxidase | PF01593 | 0.0 | Low |
| 28. TraesCS5B02G049800 | Protoporphyrinogen oxidase | PF01593 | 0.0 | Low |

| | | | | |
|---|---|---|---|---|
| 29. TraesCS7D02G548100 | NBS-LRR disease resistance protein | PF00931 | 0.00 | Low |
| 30. TraesCS1D02G231700 | Transducin/WD40 repeat-like superfamily protein | PF00400 | No | No |
| 31. TraesCS1D02G340500 | Gamma-glutamyl hydrolase | PF07722 | No | No |
| 32. TraesCS2A02G064600 | PHD finger protein | | No | No |
| 33. TraesCS2A02G564400 | Harpin-induced-like protein | | No | No |
| 34. TraesCS2A02G586900 | Mevalonate kinase | PF00288; PF08544 | No | No |
| 35. TraesCS3A02G167000 | S-type anion channel | PF03595 | no | No |
| 36. TraesCS4A02G344700 | Leucine-rich repeat receptor protein kinase family protein | PF08263; PF13855; PF00560 | No | No |
| 37. TraesCS5A02G301800 | L-threonine 3-dehydrogenase | | No | No |
| 38. TraesCS5A02G467700 | RING/U-box superfamily protein | PF14369; PF13639 | no | no |
| 39. TraesCS5D02G526800 | Leucine-rich repeat receptor-like protein kinase family | PF13855 | no | No |
| 40. TraesCS6A02G312100 | ATP-dependent RNA helicase family protein | PF00270; PF00271; PF08148 | no | No |
| 41. TraesCS6B02G314800 | Enhancer of mRNA-decapping protein 4 | PF16529 | No | No |
| 42. TraesCS6D02G266500 | Enhancer of mRNA-decapping protein 4 | PF16529 | No | No |
| 43. TraesCS7A02G429300 | T-box transcription factor, putative (DUF863) | | No | No |
| 44. TraesCS7A02G538500 | Glyoxylate reductase/hydroxypyruvate reductase | PF00389; PF02826 | No | No |
| **5'UTR IRE Motif** | | | | |
| 1. TraesCS2A02G459700 | DNA mismatch repair protein MutL | | -11.30 | High |
| 2. TraesCS6D02G402800 | Zinc finger CCCH domain-containing protein 19 | PF15663 | -5.70 | High |
| 3. TraesCS5A02G147500 | Remorin | PF03763 | -5.20 | High |
| 4. TraesCS7A02G006900 | Disease resistance protein RPP13 | PF00931 | -5.10 | High |
| 5. TraesCS7D02G399600 | Cupredoxin superfamily protein, putative | | -10.10 | Medium |
| 6. TraesCS5D02G219100 | Importin subunit beta-1 | PF03810; PF13513 | -7.00 | Medium |
| 7. TraesCSU02G028000 | F-box domain containing protein, expressed | | -6.90 | Medium |
| 8. TraesCS3D02G038300 | Haloacid dehalogenase-like hydrolase superfamily protein | PF13419: | -6.70 | Medium |
| 9. TraesCS3D02G274900 (No predicted 5' UTR, Checked on cDNA Transcript) | Alpha-glucosidase | PF16863; PF13802; PF01055 | -6.40 | Medium |
| 10. TraesCS6B02G148400 | Disease resistance protein (NBS-LRR class) family | PF00931 | -5.50 | Medium |
| 11. TraesCS1B02G226000 | receptor kinase 1 | PF08276; PF07714 | -5.00 | Medium |
| 12. TraesCS5D02G481800 | Chlorophyll a-b binding protein, chloroplastic | PF00504: | -4.10 | Medium |
| 13. TraesCS7D02G286500 | Hexosyltransferase | PF13334; PF01762 | -3.30 | Medium |
| 14. TraesCS4B02G010400 | Golgin candidate 1 | PF09787 | -2.60 | Medium |

| | | | | |
|---|---|---|---|---|
| 15. TraesCS2D02G407000 | Target of rapamycin complex 2 subunit MAPKAP1 | | -0.20 | Medium |
| 16. TraesCS5D02G094600 | Proton pump interactor, putative | | 0.00 | Medium |
| 17. TraesCS1B02G110600 | Pectin lyase-like superfamily protein | PF00295 | No Result | No Result |
| 18. TraesCS1D02G078700 | Expansin protein | PF03330; PF01357 | No Result | No Result |
| 19. TraesCS2D02G289800 | NAC domain protein | PF02365 | No Result | No Result |
| 20. TraesCS5A02G040500 | Aldo-keto reductase, putative | PF00248 | No Result | No Result |
| 21. TraesCS5A02G186800 | Kinase, putative | PF00069 | No Result | No Result |
| **FPN IRE Motif (From Full Length cDNA Transcript)** | | | | |
| 1. TraesCS4A02G085200 | Solute carrier family 40 protein | PF06963: Ferroportin1 (FPN1) | No Result | No Result |
| 2. TraesCS4B02G219000 | Solute carrier family 40 protein | PF06963: Ferroportin1 (FPN1) | No Result | No Result |
| 3. TraesCS4B02G326200 | Solute carrier family 40 member 1 | PF06963: Ferroportin1 (FPN1) | No Result | No Result |
| 4. TraesCS4D02G219200 | Solute carrier family 40 member 3 | PF06963: Ferroportin1 (FPN1) | No Result | No Result |
| 5. TraesCS4D02G323100 | Solute carrier family 40 member 1 | PF06963: Ferroportin1 (FPN1) | No Result | No Result |
| 6. TraesCS5A02G087700 | Solute carrier family 40 protein | PF06963: Ferroportin1 (FPN1) | No Result | No Result |
| 7. TraesCS5A02G498400 | Solute carrier family 40 member 1 | PF06963: Ferroportin1 (FPN1) | No Result | No Result |
| 8. TraesCS5B02G093600 | Solute carrier family 40 protein | PF06963: Ferroportin1 (FPN1) | No Result | No Result |
| 9. TraesCS5D02G098400 | Solute carrier family 40 protein | PF06963: Ferroportin1 (FPN1) | No Result | No Result |
| 10. TraesCS7A02G357400 | Solute carrier family 40 member 1 | PF06963: Ferroportin1 (FPN1) | No Result | No Result |
| 11. TraesCS7B02G270400 | Solute carrier family 40 member 1 | PF06963: Ferroportin1 (FPN1) | No Result | No Result |
| 12. TraesCS7D02G365300 | Solute carrier family 40 member 1 | PF06963: Ferroportin1 (FPN1) | No Result | No Result |

**Table S3: UTR sequences of wheat genes showing high stability and presence of the IRE elements as represented in Table S3.** Yellow indicates the potential IRE sites, and the red-colored bases are the primers used to clone the fragment.

| | |
|---|---|
| >TraesCS2A02G206700.1 utr3:protein_coding | ACAGGGCATGCAGTGTGGATTAAGCTAAGCTTGTTGATAGAGTAGTACTATTTTGTAGGAGCTGGTTGGTTGGTGTTAGGGATTAGCTTCCTTTTTGGCTTGCACAGTGTTGCAGATCGAATTAGCGGATTGCACCGGTGGGTCTCCAGTATCCCCACCCCATGCTTCTATCCTTGGTTTATTACCCTGCTATTTTGGTGAAGTGGTCGTAACTTGTTCAAGCCGAAACTATTTCTGTTGGCATGGGCACAAGCAACGAAAAGCAAGAAAAATGAGCCGCGGGTATGATGAATCTGTTGAAAGGAAGAGTCGGTTTTGGTTCAGAATAGCT |
| >TraesCS2B02G233900.1 utr3:protein_coding | ACAGGGCATGCAGTGTGGATTAAGCTAAGCTTGTTGATAGAGTAGTACTCATTTTGTAGGAACTGGTTGGTTGGTGTTAGGGATTGTAGCTTCCTTTTTGCTTGCACAGTGTTGCAAATCAAATTAGCGGATTGCACCGGTGGATCTCTATTAT |
| >TraesCS2B02G186800.1 utr3:protein_coding | AGTAGCTCTTCGCTCTTTCTTGATTCAGATTAGCCACTGCCACTGGTGCGTTGCCGTTCGGAATGGAAGAACCTAAATGTTCCAAGTCTCACCGCTCGACAAACTATGCAGGCAATTTTCCATAAGTTGTTTCTCTAGTTTTAAATACTAATGACAGCCAGCCTCCCTGTTCCCTGCCCCATTCCGTTTTGATGATGTGTATCTAATGTGTGCGACATTCGTATAATCAGCTCCCCCTTTTTAGCAAGTTACTAGAAGCCGAGAACGGCTGTAGTGCTCAGATTGTTCTTGTTCTGAGTTGGAGAAGCAAAAAAATGCTGGTTACTCTTTTCAGTGGCAAACAAACGTTGCCAGTGAGCAATAATTTTTGAAGGGAAGGAACCACATTTGAGTATTCCGTCGATAAATTTGCAATTAAGCTAGCGTTTGTTTGGTGAGTGC |
| >TraesCS6D02G402800.1 utr5:protein_coding | GCACAGACAGCCAGAGCCAGTGAGCTCTCACCTTGGTTGGGGTGGTAATAATAAATCCAGGTCCAGGTCCAGGTCCAGGTCCAGGTCCATCCCTCTGCTTCTGCCCTCGACCCGGCAGCTCCACCTCGCCCGCGGCG |
